# N-terminal extensions strengthen hydrophobic inter-subunit interactions between HU’s C-terminal domains to frustrate heterodimer formation

**DOI:** 10.1101/2021.02.16.431378

**Authors:** Kanika Arora, Bhishem Thakur, Arpita Mrigwani, Purnananda Guptasarma

## Abstract

HU is a nucleoid-associated protein (NAP) that helps bacterial chromosomal DNA to remain compact. *Escherichia coli* contains two homologs of HU that are ~ 70 % identical: HU-A and HU-B. The early log phase, late log phase, and stationary phase of *E. coli* growth are reported to be dominated, respectively, by HU-AA homodimers, HU-BB homo-dimers, and HU-AB heterodimers. Here, we show that the formation of HU-AB heterodimers occurs to a much lower degree in HU chains that have a displaced N-terminus, whether through addition of an N-terminal affinity (polyhistidine) tag, or fusion of a fluorescent protein. A combination of mass spectrometry, spectroscopy, chromatography, and electrophoresis (exploring glutaraldehyde crosslinking of subunits) was used to study the mixing, co-expression, unfolding and refolding of HU-AA and HU-BB homodimers. The data suggests that, in HU polypeptides with N-terminal extension, whereas inter-subunit contacts between the alpha helical N-terminal domains (NTDs) undergo facile unfolding and dissociation, inter-subunit contacts between the beta sheet- and IDR-dominated C-terminal domains (CTDs) fail to do so, due to persistence of hydrophobic inter-subunit interactions between two beta sheets. This persistence causes HU to remain nominally dimeric even after substantive unfolding, and frustrates subunit exchange and heterodimer formation.

## INTRODUCTION

### HU is a nucleoid associated protein (NAP) that exists as two homologs in Escherichia coli

The genome of *Escherichia coli* contains ~4.6 million base-pairs of double-helical circular DNA (with a total circumferential length of ~1.6 mm), packed into a rod-shaped cell of 1 μm length. The enormous compaction of DNA that is required to achieve the packing of such a large amount of DNA within such a small cell is wrought by a combination of negative supercoiling of DNA, and its stabilization through binding of nucleoid associated proteins (NAPs).^1,2^ One of the twelve known types of bacterial NAPs is HU. Whereas in most bacteria, there is only one form (or homolog), in *Enterobacteriaceae* like *E. coli*, HU exists as two homologs, known as HU-A and HU-B, produced by two closely-related genes, known as *hupA* (which produces HU-A), and *hupB* (which produces HU-B), which are located at 90.5 minutes, and 9.9 minutes, respectively, on the *E. coli* chromosome.^3,4^

### HU is a highly-abundant protein that is predominantly a homo/hetero dimer, with minority populations of higher order associations

HU sometimes exists as a homodimer. The homodimers of HU-A, and HU-B, respectively, are HU-AA and HU-BB. Sometimes, HU exists as a heterodimer, with HU-A and HU-B chains associating to generate HU-AB. A fraction of every HU population is associated into higher order forms such as tetramers and octamers.^5,6^ In the cell, the highest concentration of HU is reached during the late exponential phase of growth, and in the stationary phase, rising up to 50,000 homo/hetero dimers of HU per cell.^7,8^ This allows us to calculate the yield of naturally-produced HU from a culture. Since cultures in the stationary phase commonly have an optical density of over 2.0 at 600 nm, and since each unit of optical density at this wavelength amounts to at least 8×10^8^ cells per ml (or 1.6 × 10^9^ cells per ml of culture, each containing ~ 5×10^4^ dimers of HU), it can be seen that a single liter of *E.coli* culture in the stationary phase contains over 2.5 mg of HU, even without any deliberate overexpression of either HU-A or HU-B chains from *hupA* and *hupB* genes mounted on multi-copy plasmids. Therefore, HU may be thought of as a highly abundant protein.

### The concentrations of HU-AA, HU-BB and HU-AB vary during growth

In *E. coli*, the cellular concentrations of different HU homodimers and heterodimers vary with the phase of bacterial growth,^7^ owing to differential activities of the promoters of *hupA* and *hupB*.^6^ HU-A polypeptides are expressed from a single promoter on the *hupA* gene, while HU-B polypeptides are expressed from three different promoters (P2, P3 and P4) on the *hupB* gene. The *hupA* gene is transcribed at high frequency during early stages of exponential growth, causing the homodimer, HU-AA, to dominate the cellular HU population. In contrast, the *hupB* gene is transcribed more frequently from promoter P2 during the late exponential phase, and from promoter P3 during the transition to stationary phase,^6^ causing the homodimer, HU-BB, to dominate the late exponential phase, and the heterodimer, HU-AB, to dominate the stationary phase.^5,6^ Importantly, the expression of both HU-A and HU-B polypeptide chains continues throughout all phases of growth,^6^ with only the relative amount of the two chains varying, and leading to changes in the populations of homo/hetero dimers.

### HU binds non-specifically to different forms of DNA and induces DNA bending

HU binds to DNA in a non-sequence specific manner, and shows higher affinity for bent DNA, nicked DNA, gapped DNA, and cruciform (four-way junction) DNA (4WJ-DNA) than standard B-form DNA.^9.10^ Dimers of HU mainly bind to DNA through two extended beta hairpin loops, one of which happens to be derived from each monomer. These lysine- and arginine-rich beta hairpin loops embrace the minor groove of DNA from opposite sides, with the two subunits crossing the C-terminal regions of their chains much like a pair of scissor tongs.^11^ The two antiparallel beta sheets that anchor the DNA-binding beta hairpin loops also themselves contact (i.e., abut) the major groove of DNA.

Further, HU possesses additional DNA-binding sites, e.g., there is a second DNA-binding site that consists of a cluster of two positively charged residues, K83 and K86, located in a short helix present at the C-terminus of each monomer.^12.13^ There are also other positively charged residues that could potentially contact DNA, and especially DNA that is wrapped around the HU dimer, and such residues could include the positive charged residues that are present near the N-terminus, or the alpha amino group present at the N-terminus itself, with some proximal positively-charged residues. Binding of HU induces bending of DNA. Such bending is presumed to assist in the compaction of DNA by HU.^11^ As already mentioned, HU also shows high affinity for DNA that is bent in any fashion.

### Conflicting reports about the structural contents and stabilities of HU-AA and HU-BB homodimers

During the thermal unfolding of ion exchange-purified HU-AA or HU-BB homodimers, the dimeric state (called N2) is reported to first unfold into partially unfolded dimeric intermediates (called I2) that subsequently transform into fully unfolded and dissociated monomers (called 2D).^14^ It has been proposed that I2 is the preferred state of HU-BB, involving retention of HU-BB in a partially unfolded state at all times, with this state also being relevant to formation of heterodimers through facile subunit exchange. However, this proposal seems to be contrary to other reports which indicate that HU-BB displays a greater tendency to form higher-order oligomers than HU-AA;^5^ this is surprising, since partial unfolding is more likely to be detrimental to adoption of multimeric states, than supportive of the adoption of such states.

Evidently, not enough is understood about HU-AA and HU-BB. In fact, it is not even clear why *E. coli* has two homologs, since deletion mutants of *hupA* as well as *hupB* display only subtle growth defects, with only double mutants (involving deletion of both *hupA* and *hupB*) displaying profound defects in growth and division.^15^ Interestingly, deletion of both *hupA* and *hupB* is not lethal, although it leads to multiple defects.^16^ This indicates that HU is not an essential protein for the survival of *E. coli*. Thus, the existence of two homologs, and their complex arrangements for expression through multiple promoters, in varying amounts during the growth of the population, appears to be even more interesting from an evolutionary viewpoint. It would appear that HU-A and HU-B offer survival benefits without being essential for survival.

### HU-A and HU-B are similar but not identical polypeptides, and prefer to form heterodimers

HU-A and HU-B polypeptides display high sequence identity (~70%), high sequence similarity (~ 80%),^17^ and high structural similarity (i.e., a backbone trajectory superimposition of 0.56 Å RMSD between the structures of the HU-A monomer, 1MUL,^14^ and HU-B monomer, 4P3B, as calculated by the software, TM-ALIGN^18^). The structural similarity between an HU-A monomer (Figure 1A) and a HU-B monomer (Figure 1B) is evident from Figure 1C in which HU-A and HU-B chains are shown superimposed by the software TM-ALIGN. From Figure 1C, it is evident that there is significant conservation of both backbone trajectory and hydrophobic side-chain orientations between the HU-A and HU-B homologs. Incidentally, HU-A and HU-B also display entirely similar modes of DNA binding, although they show different substrate specificities in binding of different types of DNA,^17^ potentially owing to HU-BB homodimers displaying a greater tendency to associate to form tetramers and octamers, than HU-AA homodimers. As already mentioned, HU also forms HU-AB heterodimers that dominate the genome in the stationary phase.^5^

**Figure 1:**
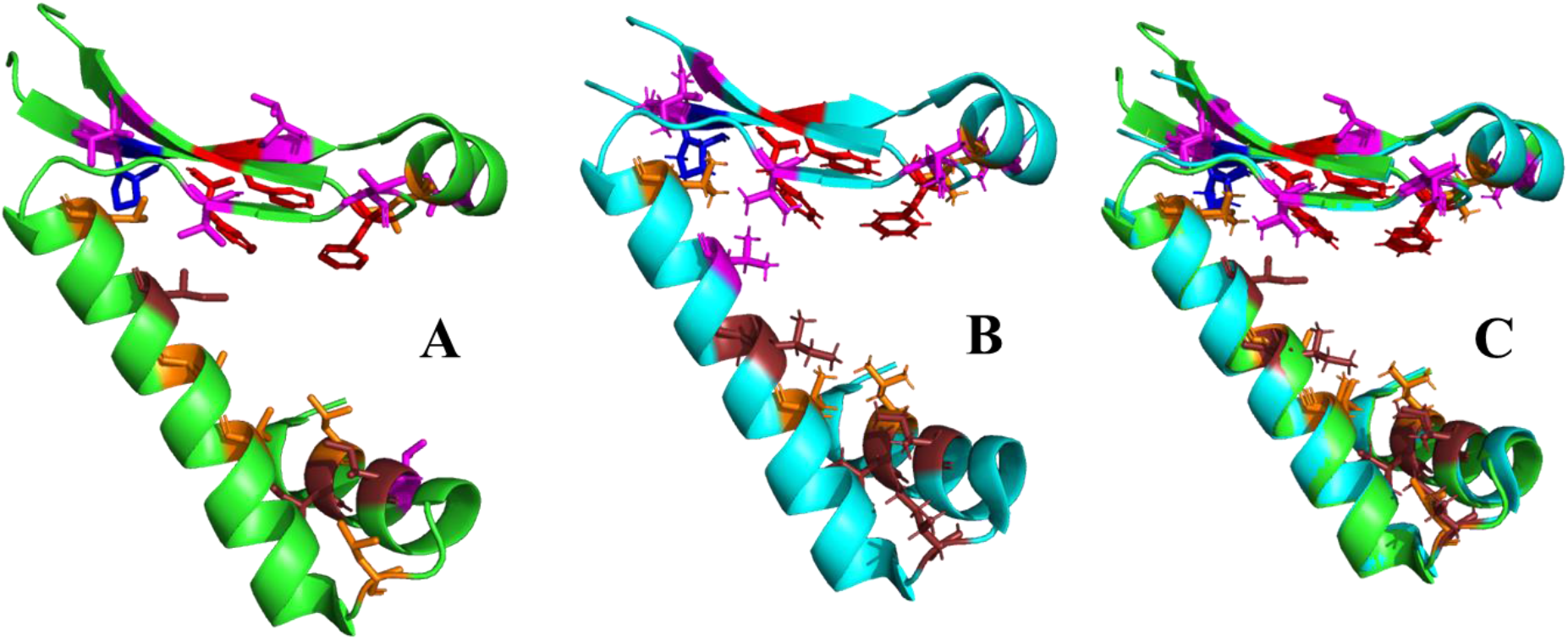
Structural similarities of *E. coli* HU homologs. *Panel A.* Structure of HU-A (PDB ID:1MUL) showing side-chains of hydrophobic residues, phenylalanine (red), proline (blue), valine (magenta), leucine (orange), and isoleucine (copper). *Panel B.* Structure of HU-B (PDB ID: 4P3V) showing hydrophobic residues with the same color scheme as in Panel A. *Panel C.* Superimposed structures of HU-A and HU-B showing conservation of backbone trajectory and hydrophobic side-chain orientations.

### The structure of HU consists of two structurally distinct domains: The helical N-terminal domain (NTD) and the sheet- and IDR-dominated C-terminal domain (CTD)

As is evident from Figure 1, in both forms of HU the polypeptide chain consists of an N-terminal domain (NTD) that contains only alpha helices, and a C-terminal domain (CTD) that consists of a network of antiparallel beta strands and disordered regions. Both NTDs and CTDs engage in inter-subunit interactions in dimers, with the NTD contacting the NTD, and the CTD contacting the CTD. However, the CTD-CTD interactions between the antiparallel beta strands are predominantly hydrophobic in nature, in comparison with interactions between the helices of the NTDs which are both ionic and hydrophobic, with the former being more important than the latter. Evidence of a degree of autonomy in HU’s CTD has been recently established by a paper from our group.^19^ In this paper, a simulacrum of the DNA-binding function of HU was successfully generated by fusing the CTDs of two different HU polypeptides through a linker peptide, to generate a novel nature-inspired DNA-binding protein, called HU-Simul. No experiment has yet probed the autonomy of the NTDs. Moreover, there remains some confusion in the literature about the status of HU-BB dimers. Some authors think that HU-BB forms oligomers more readily that HU-AA dimers.^5^ Others think that HU-BB remains largely unstructured and ready to engage in subunit exchange to form heterodimers.^14^

### HU-AB heterodimers are assumed to arise through facile subunit exchange

HU-AB heterodimers are assumed to arise through subunit exchange between HU-AA and HU-BB homodimers. It is reported that when both HU-A and HU-B chains are present together in any solution in the form of HU-AA and HU-BB homodimers, subunit exchange causes the dominant population in the mixture to eventually consist of 90 % HU-AB heterodimers, with only minority populations of HU-AA and HU-BB also present.^5,6,20^ This suggests that HU-AB is more stably formed than either HU-A or HU-B. It is also reported that HU purified from stationary phase *E. coli* consists almost exclusively of HU-AB heterodimers. Experimentally, the distinction between homodimers and heterodimers gets drawn either through separation of HU populations into monomers, followed by purification using ion exchange chromatography (which separates HU-A and HU-B), or through analytical electrophoretic separation of HU-A and HU-B polypeptides on gels including urea and Triton X-100 gels.^20^ In principle, other analytical methods such as mass spectrometry and analytical gel filtration chromatography could also be used, especially with differential mass-tagging of HU-A and HU-B polypeptides through genetic fusions with other peptides/polypeptides (e.g., fluorescent proteins).

### Overexpressed N-terminally histidine-tagged HU is reported to behave like wild-type HU in every respect

HU has been overexpressed both without, and with a 6xHistidine tag at its N-terminus in *E. coli.*^20^ HU produced with an N-terminal histidine tag (whether HU-A or HU-B) is reported to be identical to wild-type HU (lacking the histidine tag) in respect of both its structural features and its binding of DNA. Each homolog of such histidine-tagged HU (i.e., whether HU-A or HU-B) is found to be capable of compensating for the subtle growth defects shown by single-gene deletion mutants of the wild-type form of the same homolog, through complementation. Each homolog also compensates substantially for the more profound growth defects shown by double-gene deletion mutants lacking both homologs.

Moreover, when histidine-tagged HU-A and HU-B are produced, purified, completely-unfolded, mixed in equal proportions and allowed to refold, it is reported that 90 % of the resultant population is heterodimeric,^20^ exactly as observed with mixing of wild-type protein homologs that have separately purified through ion-exchange chromatography, before mixing and without first being caused to undergo unfolding and refolding.^5^ Notably, there is no separate comment about the behavior of histidine-tagged HU in this regard, i.e., about whether mixing of pre-folded N-terminally histidine-tagged HU-A and HU-B populations leads to formation of heterodimers.

Thus, if we take all known observations into account, i.e., (a) that mixing of pre-folded HU-AA and HU-BB results in formation of ~ 90 % HU-AB,^5,6^ and (b) that co-refolding of either wild-type HU-A and HU-B, or 6xHis tagged HU-A and HU-B, also leads to ~ 90 % heterodimers,^20^ and neglect (c) the lack of data about the consequences of mixing pre-folded N-terminally histidine-tagged HU-A and HU-B, it may be concluded that HU-A and HU-B polypeptides prefer to exist as heterodimers, rather than as homodimers. In other words, only residual or leftover populations of HU-A or HU-B chains (that are unable to form heterodimers, due to paucity of the other chain when there is a molar excess of one chain over another) are expected to form HU-AA or HU-BB homodimers, whenever both HU-A and HU-B polypeptides are present together in the same solution.

### Unexpectedly, affinity-purified N-terminally histidine-tagged HU homologs are purified as homodimers

Intriguingly, although this has not been remarked upon (or pointed out) previously, in this context, affinity-purified HU-A and HU-B polypeptides that are 6xHis tagged at their N-termini are reported to be homodimers of HU-AA and HU-BB,^20^ rather than heterodimers. The authors who reported this appear to have assumed that homodimer formation by the HU homolog owes to the greater presence in the cell of the same HU homolog on account of overexpression. However, we find this to be surprising for the following reason.

The studies of such histidine-tagged HU homologs report only moderate levels of overexpression of ~10 mg per liter of culture, for either of the histidine-tagged HU homologs. As already remarked, the natural level of wild-type HU in stationary phase *E. coli* is ~ 2.5 mg per liter of culture. Thus, the purification of homodimers of any HU homolog expressed at a level of ~ 10 mg per liter of culture is unexpected (where the anticipation is that heterodimers must form), because there is only a four-fold difference in the levels of the overexpressed histidine-tagged HU homolog (10 mg per liter) and the wild type HU-A and HU-B polypeptides (2.5 mg per liter). Ordinarily, with such a small four-fold difference in levels, there would be abundant scope for association between wild-type HU homolog chains and 6xHis tagged HU homolog chains of the opposite variety (i.e., HU-A with HU-B, or *vice versa*), if there were indeed facile subunit exchange occurring in both populations. Therefore, the harvesting of pure homodimers of a single overexpressed N-terminally histidine-tagged HU homolog (rather than a mixture of heterodimers and homodimers) suggests that histidine-tagged HU behaves differently, and does not display facile subunit exchange.

### Scope of the present work

Since there are no previous reports of any issues with the dissociation of wild-type HU homo/hetero dimers, and since histidine-tagged HU has not been examined in any detail thus far from a structural-biochemical viewpoint, we decided to go ahead and examine whether dimers of histidine-tagged HU homologs display problems with dissociation of subunits, or with the association of such subunits with other forms of HU (e.g., with wild-type HU, or HU fused with other proteins, such as fluorescent proteins). We also decided to examine the abilities of pre-folded and *de novo* co-expressed histidine-tagged HU homologs to form heterodimers, through (i) mixing of pre-folded populations of labelled HU-homologs, or (ii) through co-expression of a histidine tagged homolog with a non-histidine-tagged homolog of the opposite nature. We report that histidine-tagged HU has difficulty in unfolding completely and, therefore, in undergoing facile subunit exchange, ostensibly due to the relocation of the N-terminus and the effects of such relocation upon the stability of inter-subunit (CTD-CTD) hydrophobic interactions between two beta sheets.

In principle, since HU-B chains prefer to form ~ 90 percent heterodimer populations, if the histidine-tagged HU were truly identical to wild-type HU in all respects, it would be expected to engage in exchange of subunits with wild-type HU inside the cell in which it is produced, and thus contain three forms of HU: (i) homodimers of the overexpressed histidine-tagged HU homolog, e.g., HU-A; (ii) dimers of the overexpressed histidine-tagged HU-A homolog and the wild-type (untagged) homolog of the same nature, i.e., also HU-A; and (iii) heterodimers of the overexpressed histidine-tagged HU-A homolog and the wild-type of the opposite HU homolog (also untagged), i.e., HU-B. Different analytical methods, including mass spectrometry, electrophoresis and chromatography, could then be used to distinguish between such populations of different dimers. Thus, it would be a good idea to confirm whether indeed the histidine-tagged HU homolog which is purified consists of only the first of the above three species, i.e., pure histidine-tagged homodimer, and no trace amounts of the other two species (wild-type HU-A and HU-B). This would shed light on whether there is a barrier to subunit exchange in N-terminally histidine-tagged HU; a barrier that potentially prevents it from forming heterodimers as long as it is in a pre-folded form, allowing histidine-tagged HU to form heterodimers only through either unfolding and refolding, or through fresh association during biosynthesis in the cell. To examine whether histidine-tagged HU homologs are indeed pure homodimers, we produced recombinant N-terminally 6xHis tagged forms of HU-A and HU-B, and purified these and subjected them to mass spectrometry, to detect the presence of the opposite HU homolog. Here, we show that affinity-purified HU-A and HU-B are indeed pure homodimers with no trace amounts of any other species.

Further, we report that histidine-tagged HU-A and HU-B differ substantially between themselves, in respect of the temperatures required for structure melting (T_m_), and the denaturant concentrations required for structural unfolding (C_m_), or for subunit-subunit dissociation accompanying chain unfolding. We show that the dimeric interface in histidine-tagged HU homodimers (i.e. both HU-AA and HU-BB) tends to be extremely stable to urea, with the protein chains appearing to be completely unfolded by urea at a much lower concentration (e.g., ~2 M urea) than those at which crosslinking by glutaradehyde shows them to be still nominally dimeric (e.g., ~ 4 M urea). We show that the interface between subunits is differentially stable in HU-AA and HU-BB, with HU-BB being much more stable to dissociation than HU-AA, and remaining quite stable up to ~4 M urea. We show that high concentrations (50 % v/v) of the hydrophobic solvent 1,4-dioxane break the monomer-monomer interface in HU-BB, by interfering with the hydrophobic interactions between the antiparallel beta sheets contributed to the interface by each monomer, with the applicant of this solvent alone (in the absence of urea) also dissociating HU-BB dimers into monomers at solvent concentrations similar to those at which it the solvent works alone to achieve such dissociation.

This indicates that the breaking of (a) hydrogen bonds by urea which leads to the unfolding of the helices in the NTDs of HU, as well as the breaking of (b) hydrogen bonds that could be presumed to destabilize the beta sheets of the CTDs, together have insufficient potential to dissociate HU because they fail to destabilize the hydrophobic interactions between the beta sheets contributed by each monomer to the CTD-CTD interface. The thermal and chemical denaturation studies together show that the application of denaturants does not completely abolish the secondary structural content of HU, with residual signal in the region of the CD spectrum that is characteristic of beta sheets. Furthermore, we also show that the thermally-unfolded and refolded HU dimers (both HU-AA, and HU-BB) possess the ability to bind to DNA (4WJ-DNA) after refolding, and possess the same hydrodynamic volume as the wild type HU homodimers. Similarly, we show that the HU dimers unfold by dioxane, urea, or a combination of dioxane and urea are also able to refold into dimers that display crosslinking by glutaraldehyde. Finally, we show that co-expression of non-affinity-tagged RFP-HU-A and affinity-tagged (6xHis-tagged) HU-B from a single plasmid that expresses both chains simultaneously, produces a mixture of HU-BB homodimers and HU-AB heterodimers after purification using the affinity tag.

Thus, we conclude that for histidine-tagged HU homologs formation of heterodimers via subunit exchange is more improbable than formation of heterodimers through *de novo* synthesis of chains and their association, because one chains have formed a dimer it is very difficult to separate them fully. The rationale behind the complete study is explained in the figure 2.

**Figure 2:**
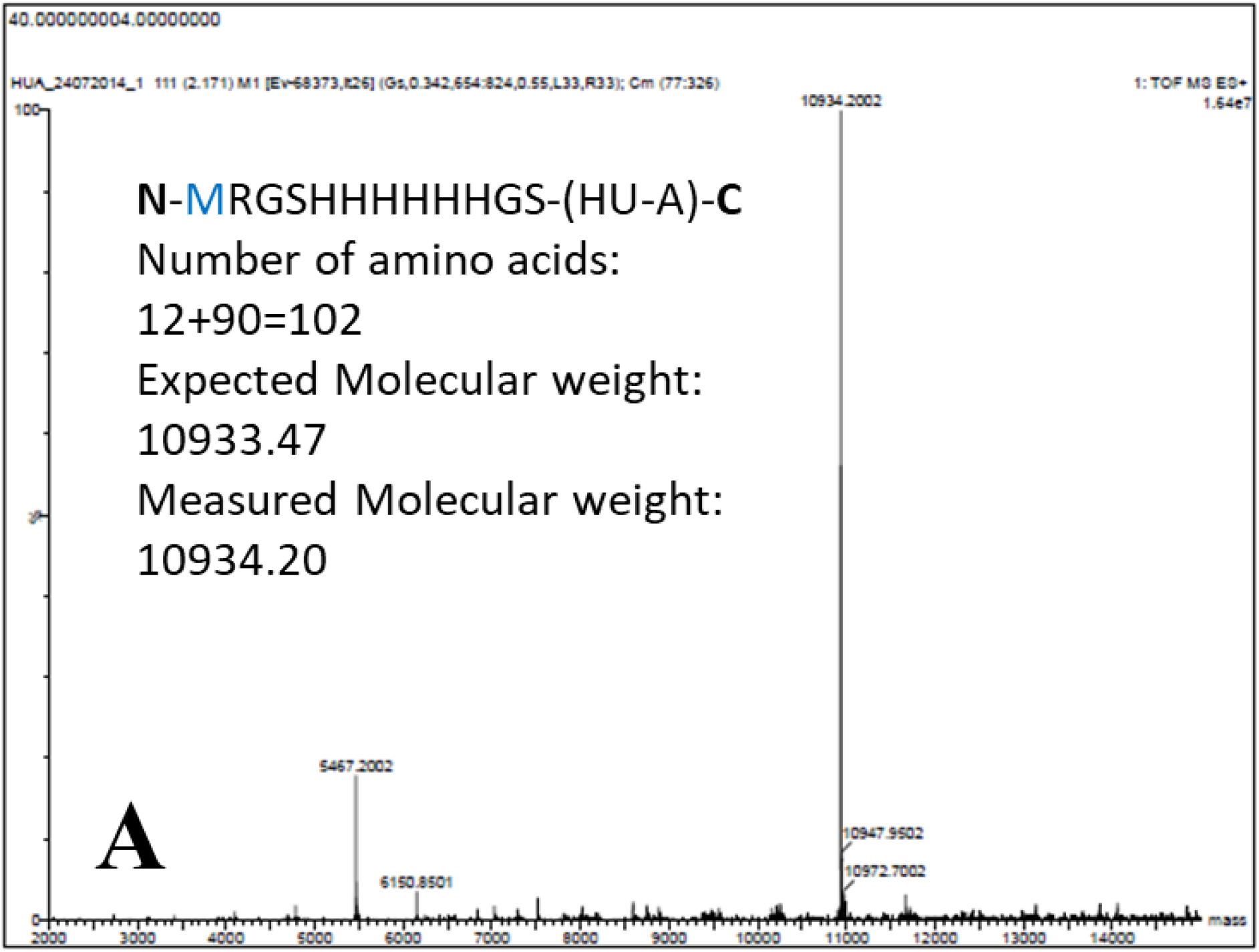
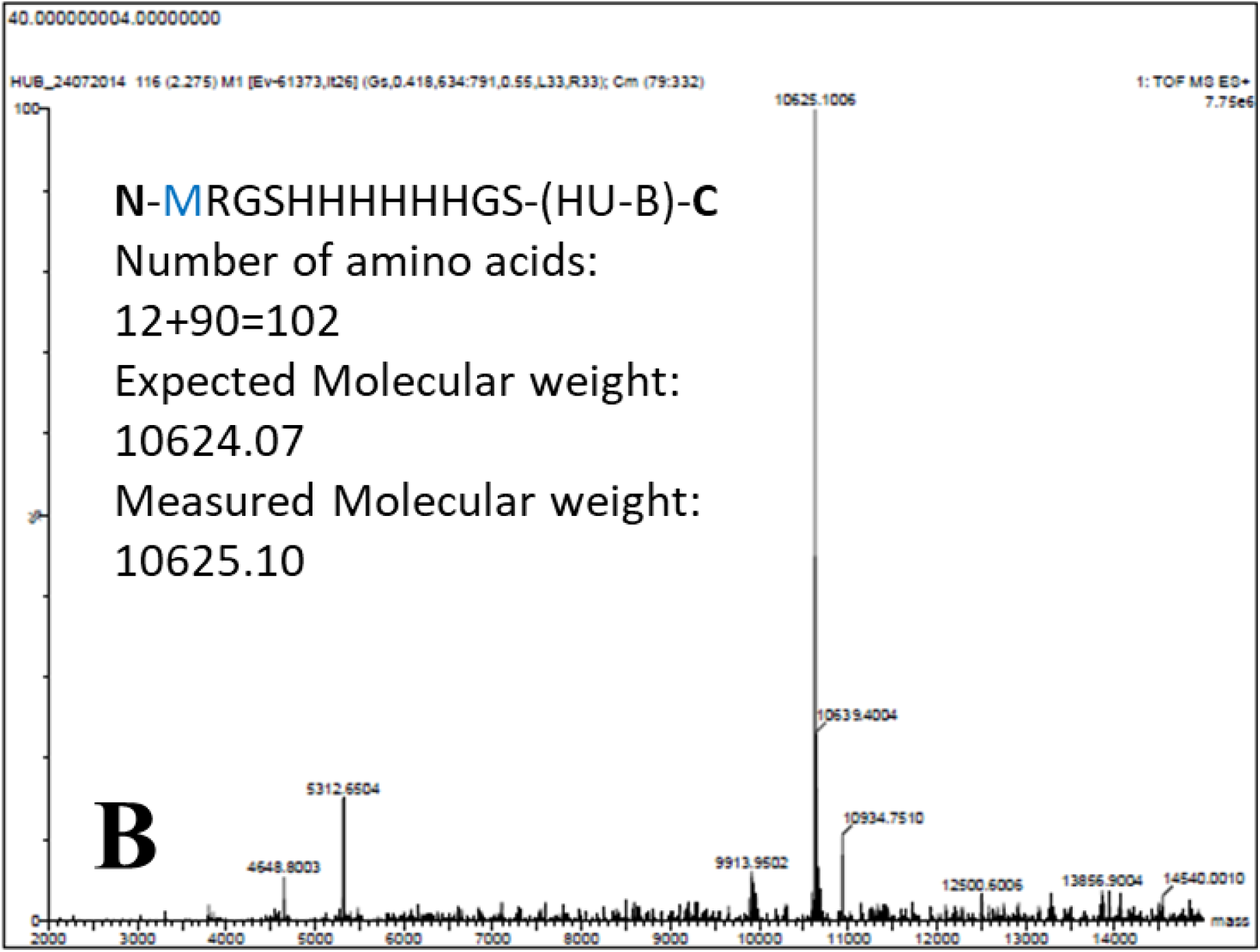
Non-histidine tagged HU is not co-purified with histidine-tagged HU. *Panel A.* MALDI-Q-TOF MS spectrum of histidine-tagged *E. coli* HU-A. The sequence of the histidine tag is shown in an inset, along with values of expected and measured molecular weights. *Panel B.* MALDI-Q-TOF MS spectrum of histidine-tagged *E. coli* HU-B. The sequence of the histidine tag (identical to that used for HU-A) is shown in an inset, along with values of expected and measured molecular weights.

## MATERIALS AND METHODS

### Cloning

(A) The *hupA* and *hupB* genes of *E.coli* K-12 sub-strain MG1655, encoding HU-A and HU-B, were cloned into the multiple cloning site (MCS) of the pQE-30 vector (Qiagen) between its Bam HI and Hind III restriction sites. Plasmids incorporating the said genes were transformed into *E.coli* M15 cells for controlled expression of 6xHis tagged versions of HU-A and HU-B, through induction by 1 mM IPTG. Further, (B) a gene encoding Tag-RFP fused to HU-A (i.e., RFP-HU-A) without a 6xHis tag was cloned between the Nde I and Xho I restriction sites of one MCS of the pET-Duet-1 plasmid vector, while the *hupB* gene encoding HU-B with an N-terminal 6xHis tag was cloned between the Bam HI and Hind III restriction sites of the second MCS in the same pET-Duet-1 vector (both sites being under T7 promoters). Following cloning, the pET-Duet-1 plasmid was transformed into BL21(DE3)pLysS* cells for controlled protein expression through induction by 1 mM IPTG. The sequences of primers used for cloning *hupA* (encoding HU-A) and *hupB* (encoding HU-B) are given below.

HU-A Bam HI-Forward: AGCTACT<u>GGATCC</u>ATGAACAAGACTCAACTGATTG
HU-A Hind III Reverse: ATATAT<u>AAGCTT</u>TTACTTAACTGCGTCTTTCAGTGCCTTG
HU-B Bam HI-Forward:AGCTACT<u>GGATCC</u>ATGAATAAATCTCAATTGATCG
HU-B Hind III Reverse: GAATACT<u>AAGCTT</u>TTAGTTTACCGCGTCTTTCAGT
HU-A Nde I Forward: AGCTACT<u>CATATG</u>AACACGACTCAACTGATTGATG
HU-A Xho I Reverse:GAATACTCT<u>CTCGAG</u>TTACTTAACTGCGTCTTTCAGTGC
HU-B Duet-1 Bam HI-Forward: AGCTACT<u>GGATCC</u>AATGAATAAATCTCAATTGATCG
HU-B Duet-1 Hind III Reverse: GAATACT<u>AAGCTT</u>TTAGTTTACCGCGTCTTTCAGT

The sequences of primers for cloning Tag-RFP and fusing it to HU-A, based on previous work,^21^ are given below.

RFP-Nde-1 Forward: 5-ATATATCATATGGCTAGCATGACTGGTGGACAGCAAATG-3

### Protein purification

After expression, cells were subjected to sedimentation, resuspension and lysis through sonication in the presence of lysozyme. HU-A or HU-B proteins were purified from the supernatants of cell lysis extracts through use of Ni-NTA affinity purification, or imidazole metal affinity chromatography (IMAC). Since HU-A and HU-B tend to be co-purified with attached fragments of DNA and other DNA-binding proteins, lysis was performed in 20 mM phosphate buffered saline (PBS) of pH 7.4, containing 1 M NaCl and 10 mM imidazole, with NaCl serving to dissociate HU from DNA. Thereafter, the extract was loaded onto the Ni-NTA column, and the column (now bound by 6xHis-tagged HU-A or HU-B) was washed with 20 mM PBS of pH 7.4, containing 2 M NaCl and 20 mM imidazole, to allow the removal of any DNA and other proteins bound to the HU polypeptides, as well as any contaminating proteins loosely bound to the column matrix. Thereafter, PBS buffer of pH 7.4 containing 250 mM imidazole was used to elute the HU polypeptides from the Ni-NTA resin. Following elution, HU polypeptides were dialysed extensively against 20 mM PBS alone, to remove imidazole, prior to spectroscopic (or other) investigations.

### Creation of the 4-way junction holiday intermediate (4WJ-DNA)

Four oligonucleotides designed to associate to create the four strands of a holiday junction intermediate were mixed together in equimolar proportions according to known protocols,^22^ heated to 90 °C and cooled from 90 °C to 25 °C over an extended period of time to allow strands to anneal into 4WJ-DNA. The sequences of the oligonucleotides used for creating 4WJ-DNA are detailed below:

Strand1: 5’-CCCTATAACCCCTGCATTGAATTCCTGTCTGATAA-3’
Strand2:5’-GTAGTCGTGATAGGTGCAGGGGTTATAGGG-3’
Strand3:5’-AACAGTAGCTCTTAATTCGAGCTCGCGCCCTATCACGACTA-3’
Strand4: 5’-TTTATCAGACTGGAATTCAAGCGCGAGCTCGAATAAGAGCTACTGT-3’

### Spectroscopic studies of chemical and thermal stability of HU-A and HU-B homodimers and their associations

A fixed concentration of HU (either 0.5 mg/ml HU-A, or HU-B) homodimers was incubated overnight with molecular biology grade urea (MP Biotech Cat No. 821530), over a range of concentrations varying from 0-8 M urea, or Guanidium Hydrochloride, or Gdm.HCl (Promega Lot No. 0000150502), over a range of concentrations varying from 0-6 M Gdm.HCl. On the following day, alterations in secondary structure owing to equilibrium unfolding were assessed through circular dichroism (CD) spectroscopy, performed using a Biologic MOS-500 CD spectrometer, and a quartz cuvette of 1 mm path length. Thermal denaturation was carried out by heating protein samples from 20 °C to 90 °C through use of the Peltier block arrangement in a Chirascan Applied Photophysics (UK) CD spectrometer, with data being collected at 2 °C intervals, using a protein concentration of 0.2 mg/ml, and cuvettes of 1 mM path length. The raw CD ellipticity data was then converted into the mean residue ellipticity (MRE) using the formula: MRE [Ɵ] in degrees = {Ɵ_observed_ (in millidegrees) x100 x MRW} /{1000 x concentration (mg/ml) x path length (cm)}. Separately, for fluorescence resonance energy transfer (FRET) fluorescence studies to examine interactions between pre-folded HU-A and HU-B, the former homolog was labelled with fluorescein isothiocyanate (FITC), and the latter homolog was labelled with tetramethylrhodamine isothiocyanate (TmRITC), with purification of labelled protein away from free label using a desalting Superdex G-25 (PD-10) column (GE Healthcare). FRET was examined using 495 excitation for FITC-HU-A, TmRITC-HU-B and mixtures of the in equimolar proportion.

### Gel filtration chromatography to establish formation of HU hetero-dimers, or examine unfolding or refolding of HU homo-dimers. Hetero-dimer formation

The eluent from the Ni-NTA purification of the expression product of the pET-Duet-1 vector in BL21(DE3)pLysS* cells was loaded onto a 24 ml Superdex-200 10/300 GL (GE Healthcare) column on an AKTA Purifier-10 workstation, to monitor the presence of Tag-RFP-HU-A along with 6xHis-tagged HU-B, by monitoring absorbance at 550 nm by RFP, and absorbance at 215 nm by HU (which naturally lacks Tyr or Trp residues and contains only three Phe residues). *Chemical unfolding of homo-dimers.* Gel filtration chromatography was used to examine the effect of urea on the hydrodynamic volumes of HU homo-dimers, by incubating protein overnight in different molar concentrations of urea in 20 mM PBS, and through loading of such protein on the same AKTA workstation using a Superdex-75 10/300 GL (GE-Healthcare) gel filtration column, except that for each urea concentration, the column was first pre-equilibrated with buffer of the same composition and urea concentration as the sample. *Refolding of homo-dimers after thermal unfolding.* HU-A and HU-B proteins (before, or after, heating at 90 °C for 5 minutes and cooling to room temperature) were loaded onto the same AKTA workstation using a 24 ml Superdex-75 10/300 GL (GE-healthcare) column mentioned above, pre-equilibrated with 20 mM PBS buffer of pH 7.4, to compare elution (and hydrodynamic) volumes of unheated and heated-cooled protein. After elution, proteins were examined for their ability to bind to 4WJ DNA using the electrophoretic mobility shift (EMSA) assay on 0.5 % agarose gels, to test heated-cooled HU protein for refolding to DNA binding-competent form. Similarly, ability of chemically-denatured HU to refold was assessed by dialyzing denaturants out and performing gel filtration chromatography to monitor the formation of dimeric HU.

### Glutaraldehyde cross-linking to examine oligomeric status during HU unfolding or refolding

The oligomeric status of HU-A or HU-B protein was estimated using protein cross-linking and SDS-PAGE analyses through use of glutaraldehyde as a chemical cross-linker present at a concentration of 0.1 % (v/v). After addition of glutaraldehyde, proteins were incubated at room temperature for 5 minutes, and 5x SDS sample loading buffer was added to an appropriate (standard) measure, followed by boiling of the sample and loading on 15 % SDS-PAGE. To examine the survival of dimeric forms, or higher-order associations, after treatment with a chemical denaturing agent capable of destabilizing hydrogen-bonding interactions, HU-A and HU-B proteins were incubated overnight in different molar concentrations of urea and subjected to chemical cross-linking and electrophoresis on SDS-PAGE, to investigate the survival of dimers or oligomers after incubation with urea. Separately, to establish the role of hydrophobic interactions in the survival of dimers of HU-A, or HU-B, proteins were incubated overnight with a nonpolar, water-miscible solvent, 1,4-dioxane (CDH fine chemicals), in the concentration range of 0-50 % dioxane (v/v) in the presence, or absence, of a fixed concentration of 4 M urea. In yet other experiments, glutaraldehyde crosslinking was conducted at 90 °C, followed by SDS-PAGE analysis, to examine persistence of dimers in HU populations at 90 °C.

## RESULTS

### Histidine-tagged HU-A and HU-B do not detectably form any heterodimers with wild-type HU-A and HU-B inside the cell

Figures 2A and 2B, respectively, show MALDI-Q-TOF MS spectra of Ni-NTA affinity chromatography purified 6xHis (histidine-tagged) HU-A and HU-B, in which the experimentally-measured masses of both species can be seen to be within a unit mass of the expected masses. Notably, neither spectrum presents any evidence for the presence of other species such as the wild-type (non-histidine-tagged) HU-A or HU-B chains, as mass peaks. Given that the presence of such chains even at trace levels would have been detectable, if present, it would appear that histidine-tagged HU-A and HU-B populations are indeed pure homodimers, as earlier reported,^20^ i.e., despite examination through a more sensitive method of detection (such as mass spectrometry), trace amounts of non-histidine-tagged HU-A or HU-B are not found. This suggests that histidine-tagged HU chains either do not dimerize with non-histidine-tagged dimers, or do so extremely poorly.

### Mixing of pre-folded histidine-tagged HU-A and HU-B homodimers yields < 5 % heterodimers, instead of ~ 90 %

Figure 3A shows the results of fluorescence resonance energy transfer (FRET) studies conducted with histidine-tagged homodimers of FITC-labelled-HU-A and TmRITC-labelled-HU-B. The experiment was conducted to examine interactions amongst the two populations through subunit exchange resulting in stable heterodimers. Such heterodimers would be expected to display FRET through reduction of FITC fluorescence and increase in TmRITC fluorescence emission. If subunit exchange occurred to a sufficient extent to generate 90 % heterodimers, substantial FRET could be expected between the FITC on HU-A and the TmRITC on HU-B. Light of 495 nm excites both FITC and TmRITC. From the emission spectra of the two populations labelled by the two fluorophores, it is evident that TmRITC-HU-B’s emission (red curve) shows almost no fluorescence signal in the peak emission range of the FITC-HU-A emission (green curve), between 500 and 550 nm, whereas FITC-HU-A’s emission (green curve) shows significant signal in the peak emission range of TmRITC-HU-B, between 550 and 600 nm. Therefore, when the green and red curves are digitally added to generate the numerical sum of the spectra, the resulting spectrum (black curve) overlaps with the green curve between 500 and 550 nm, but is distinct from both green and red curves between 550 and 600 nm, in Figure 3A.

**Figure 3:**
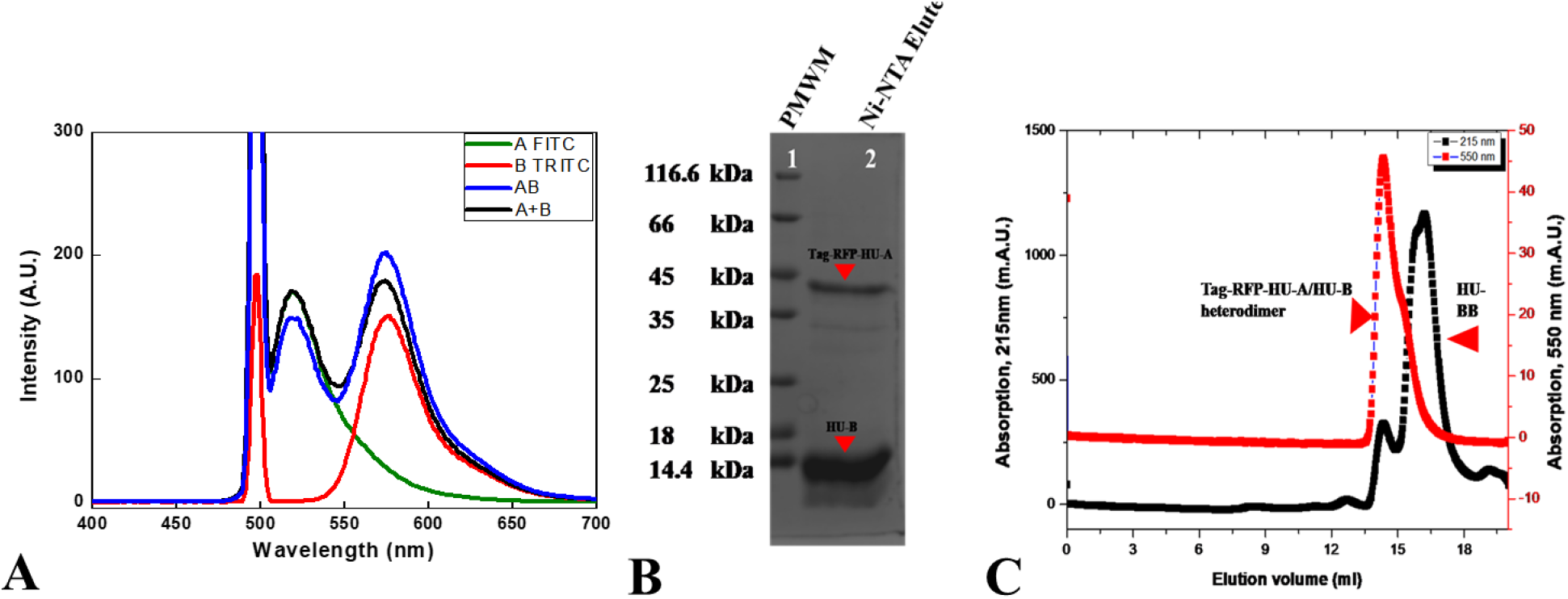
Extent of heterodimer formation obtained through mixing of HU-A and HU-B, or through co-expression. *Panel A.* Fluorescence resonance energy transfer (FRET) studies involving mixing of purified (histidine-tagged) FITC-labelled HU-A, and TmRITC-labelled HU-B, with excitation at 495 nm. Emission from the FITC on HU-A (in the absence of TmRITC-labelled HU-B) is shown in green, and from the TmRITC on HU-B (in the absence of FITC-labelled HU-A) is shown in red. The numerical sum of the green and red spectra is shown in black, as the expected spectrum in absence of FRET. The spectrum in blue shows the measured spectrum, collected to detect reduced FITC emission and increased TmRITC emission. The Rayleigh scatter peak is shown captured at 495 nm in the blue and red spectra. *Panel B.* SDS-PAGE profile of histidine-tagged HU-B purified from *E. coli* co-expressing non-histidine-tagged RFP-HU-A. *Panel C.* Gel filtration chromatographic profile of histidine-tagged HU-B purified from *E. coli* co-expressing non-histidine-tagged RFP-HU-A with monitoring of absorbance at 550 nm (RFP absorption) and 215 nm (peptide bond absorption).

The figure shows that upon mixing of equimolar amounts of the two populations, followed by incubation for 30 minutes at room temperature (which produces ~ 90 % heterodimers of wild-type non-histidine-tagged HU-A and HU-B), further followed by excitation with light of 495 nm, there is only a marginal reduction in the donor (FITC) fluorophore’s emission, along with a marginal increment in the acceptor (TmRITC) fluorophore’s emission in the experimentally-obtained spectrum (blue curve), relative to the curve obtained by digitally adding the FITC and TmRITC spectra (black curve). If the labelled histidine-tagged HU-A and HU-B populations had undergone facile subunit exchange to generate ~ 90 % heterodimers, the difference between the blue and black curves would have been profound, rather than marginal. This indicates that after mixing of the two labelled populations, heterodimers form to the extent of less than 5 %, rather than ~ 90 %. In contrast, as already noted in the introduction, mixing of pre-folded wild-type non-histidine tagged pure HU-A and HU-B homodimers (produced through unfolding of purified HU into separate chains, ion exchange chromatic separation of HU-A and HU-B chains, followed by refolding and mixing of refolded populations) yields ~ 90 % heterodimers. Evidently, therefore, histidine-tagged HU-A and HU-B do not undergo facile subunit exchange-based heterodimer formation, unlike wild-type non-histidine-tagged forms of HU that do form heterodimers.

### Co-expression of non-histidine-tagged RFP-HU-A with histidine-tagged HU-B also yields < 5 % heterodimers, instead of ~ 90 %

Figure 3B shows the analytical SDS-PAGE gel electrophoretic profile of Ni-NTA purified N-terminally histidine-tagged HU-B (~ 10.6 kDa) populations co-overexpressed with non-histidine-tagged RFP-HU-A (~ 38.6 kDa) within the same cell. Following bacterial cell lysis, during Ni-NTA-based affinity (IMAC) purification, RFP-HU-A homodimers are lost in the flow-through of the column because they have no 6xHis histidine tag. However, two other species are purified, consisting of homodimers of histidine-tagged HU-B, and heterodimers of histidine-tagged HU-B and non-histidine-tagged, but RFP-fused HU-A (RFP-HU-A). Figure 3B shows that both polypeptides (6xHis-HU-B and RFP-HU-A) are present in purified protein population, with the RFP-HU-A constituting < 10 % of the total protein purified. Assuming that the RFP-HU-A homodimers in the flow-through were equal in amount to the 6xHis-HU-B population (because both were expressed under identical promoters from the pET-Duet-1 plasmid vector, incorporating the two genes at different multiple cloning sites), this would indicate that the heterodimer population is < 5 % of the total overexpressed HU population.

Essentially, the same conclusion is reached from consideration of Figure 3C, which shows the analytical gel filtration chromatographic elution profile of the Ni-NTA (IMAC) purified protein. As already mentioned, the red coloured RFP-HU-A homodimer population is lost in the flow-through of the Ni-NTA column. What is purified, as seen in Figure 3B, is both the 6xHis-tagged HU-B homodimer and the heterodimer of 6xHis-tagged HU-B and RFP-HU-A. The gel filtration profile shows that the peak eluting at ~ 14 ml is the heterodimer of 6xHis-tagged HU-B and RFP-HU-A, because it displays absorption at 550 nm. On the other hand, the majority population of 6xHis-tagged HU-B homodimer, being smaller, elutes at ~ 16.5 ml, with no associated 550 nm absorption. Once again, from the black curve, it is evident that the peak with the associated 550 nm absorption (~ 14 ml) has < 10 % of the area of the peak without the 550 nm absorption (~ 16.5 ml). Assuming that the 6xHis-tagged HU-B homodimer and the RFP-HU-A homodimer (not captured through affinity purification) are identical in amount, it is reckoned that the heterodimer of 6xHis-tagged HU-B and RFP-HU-A is formed to a level of < 5 % of the total overexpressed HU population. This pales in comparison with reports of ~ 90 % heterodimer formation between non-histidine-tagged wild-type HU-A and HU-B,^6^ as well as histidine-tagged HU-A and HU-B, suggesting once again that histidine-tagged HU homologs are compromised in respect of heterodimer formation through subunit exchange-based mechanisms.

### Chemical denaturation reveals persistence of a hydrophobically-stabilized dimeric interface in histidine-tagged HU that could frustrate subunit exchange

We begin this part of the study by assessing the relative abilities of the 3-D structures of histidine-tagged HU-A and HU-B to withstand chemical unfolding and dissociation, using a combination of spectroscopic, electrophoretic and chromatographic experiments, with yielded the results and conclusions summarized in the following sub-sections.

#### Circular dichroism reveals differential unfolding of HU-A and HU-B by urea and Gdm.HCl, with hints of persistence of an unfolding-resistant sub-structure

Far-ultraviolet circular dicroism (CD) spectroscopy was used to assess changes in secondary structural content in HU as a function of denaturant concentration, following overnight incubation of samples of HU in denaturing buffers. For denaturation by urea, representative CD spectra are shown in Supporting Information Figures S1A and S1B. These (and the corresponding spectra for denaturation by Gdm.HCl) reveal that denaturation does not proceed to completion in a facile manner with either denaturant, i.e., either with urea, which predominantly destabilizes hydrogen bonding interactions, or with guanidium hydrochloride (Gdm.HCl), which destabilizes both electrostatic and hydrogen bonding interactions. With both denaturants, residual ellipticity is observed at 222 nm even at very high denaturant concentrations. The raw ellipticity values at 222 nm are plotted and shown in Figures 4A-4D, from which the concentrations of denaturant required to achieve fifty percent unfolding of the population (C_m_) were determined for the histidine-tagged forms of both HU-A and HU-B. For HU-A, the C_m_ was determined to be 0.82 M for denaturation by Gdm.HCl, and 1.77 M for denaturation by urea. In contrast, for HU-B, the C_m_ was determined to be 1.03 M for denaturation by Gdm.HCl, and 2.14 M for denaturation by urea.

**Figure 4:**
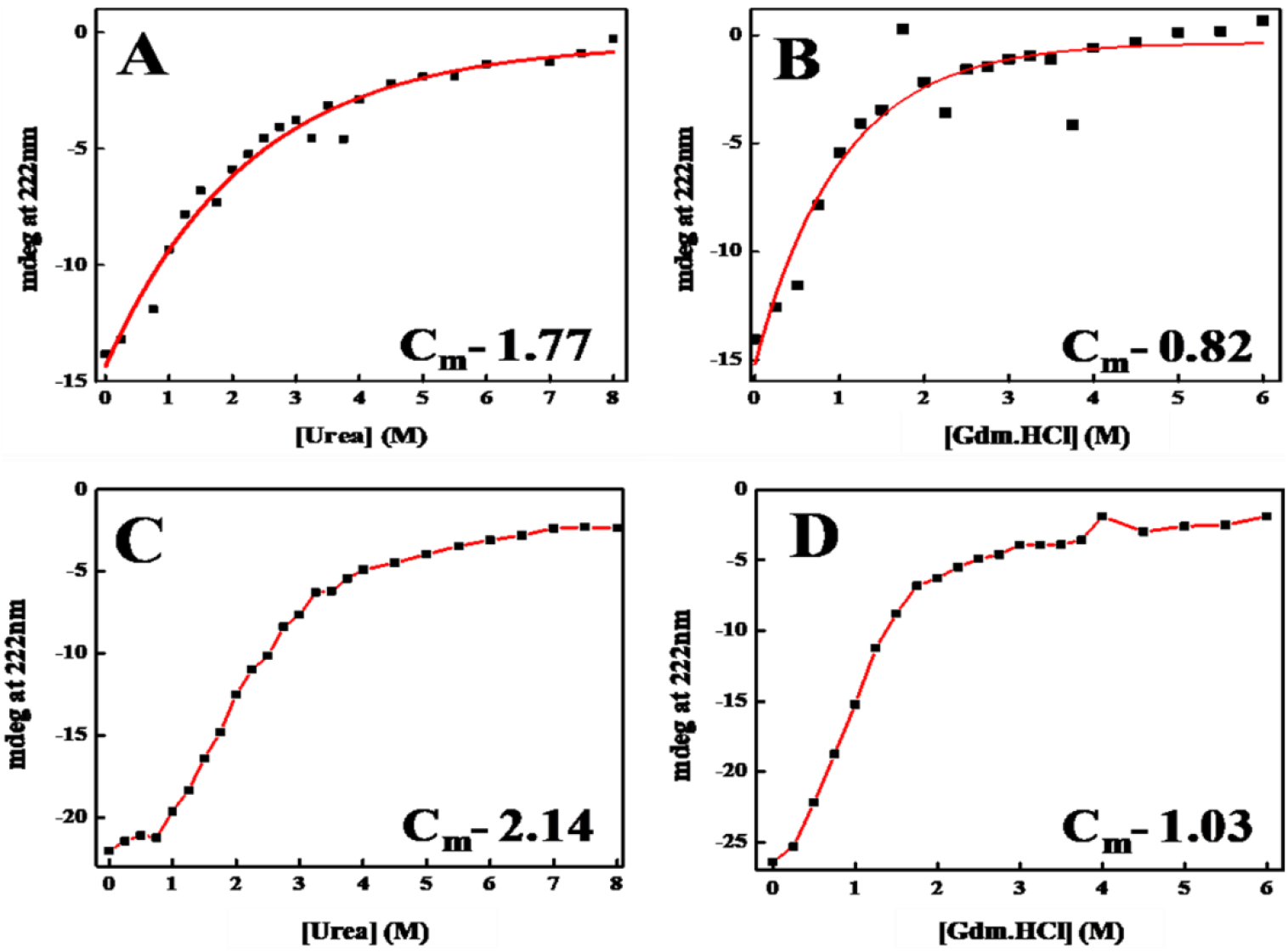
Raw circular dichroic (CD) ellipticity of HU-A and HU-B in millidegrees as a function of denaturant concentration. *Panel A.* HU-A variations between 0 and 8 M urea. *Panel B.* HU-A variations between 0 and 6 M Gdm.HCl. *Panel C.* HU-B variations between 0 and 8 M urea. *Panel D.* HU-B variations between 0 and 6 M Gdm.HCl.

Clearly, therefore, HU-B is found to be more stable to chemical denaturation than HU-A. The spectra and plots further reveal (i) that the concentration of urea required to cause unfolding is roughly twice the concentration of Gdm.HCl, for histidine-tagged forms of both HU-A and HU-B; (ii) that the histidine-tagged form of HU-A is destabilized by lower concentrations of both urea, and Gdm.HCl, than HU-B, (iii) that in both homologs, structural destabilization is initiated at extremely-low denaturant concentrations, with a very high initial rate of structural unfolding with increasing denaturant concentration (below 3 M urea, and below 1.5 M Gdm.HCl); and (iv) that the rate of unfolding is much slower at higher denaturant concentrations (above 3 M urea, and above 1.5 M Gdm.HCl).

The slowing down of the unfolding with increasing denaturant concentrations above 3 M urea, and above 1.5 M Gdm.HCl, in both HU-A and HU-B, as noted in Figures 4A-4D, and as suggested by the shapes and relative intensities of the CD spectra in Supporting Information Figures S1A and S1B, collectively suggest that both histidine-tagged homologs, HU-A and HU-B, lose their helical structures in a much more facile manner than they lose other structures (which potentially comprise a structural core). This could explain the fact that unfolding occurs in a significantly less facile manner at higher denaturant concentrations. We have not been able to find any evidence of chemical denaturation studies involving HU in the literature, prior to these studies.

#### Analytical gel-filtration chromatography reveals differential unfolding by urea of HU-A and HU-B, with hints of unfolding preceding dissociation (suggesting a persistent sub-structure)

Figures 5A and 5B, respectively, show variations in the gel filtration chromatographic behaviors of HU-A, and HU-B as a function of urea concentration, after overnight incubation in different molar concentrations of urea, and assessed after equilibrating columns with appropriate concentrations of urea. The data reveals the effect of urea upon the hydrodynamic volumes of HU dimers. With increase in urea concentration, the hydrodynamic volumes of HU samples were initially found to increase due to the formation of partially-unfolded dimers (of lower elution volume), prior to formation of dissociated and substantively-unfolded monomers (of higher elution volume). HU-A which had shifted to lower elution volumes (associated with higher hydrodynamic volumes signifying partial unfolding) at urea denaturant concentrations below ~ 3.5 M urea, was observed to shift towards higher elution volumes signifying dissociation of partially unfolded dimers into substantively-unfolded monomers. Such a shift was not seen with HU-B even at 4 M urea, supporting the result from CD spectroscopy that HU-B is more stable than HU-A. In all likelihood, the shift occurs at even higher urea concentrations that could not be accessed, for HU-B.

**Figure 5.**
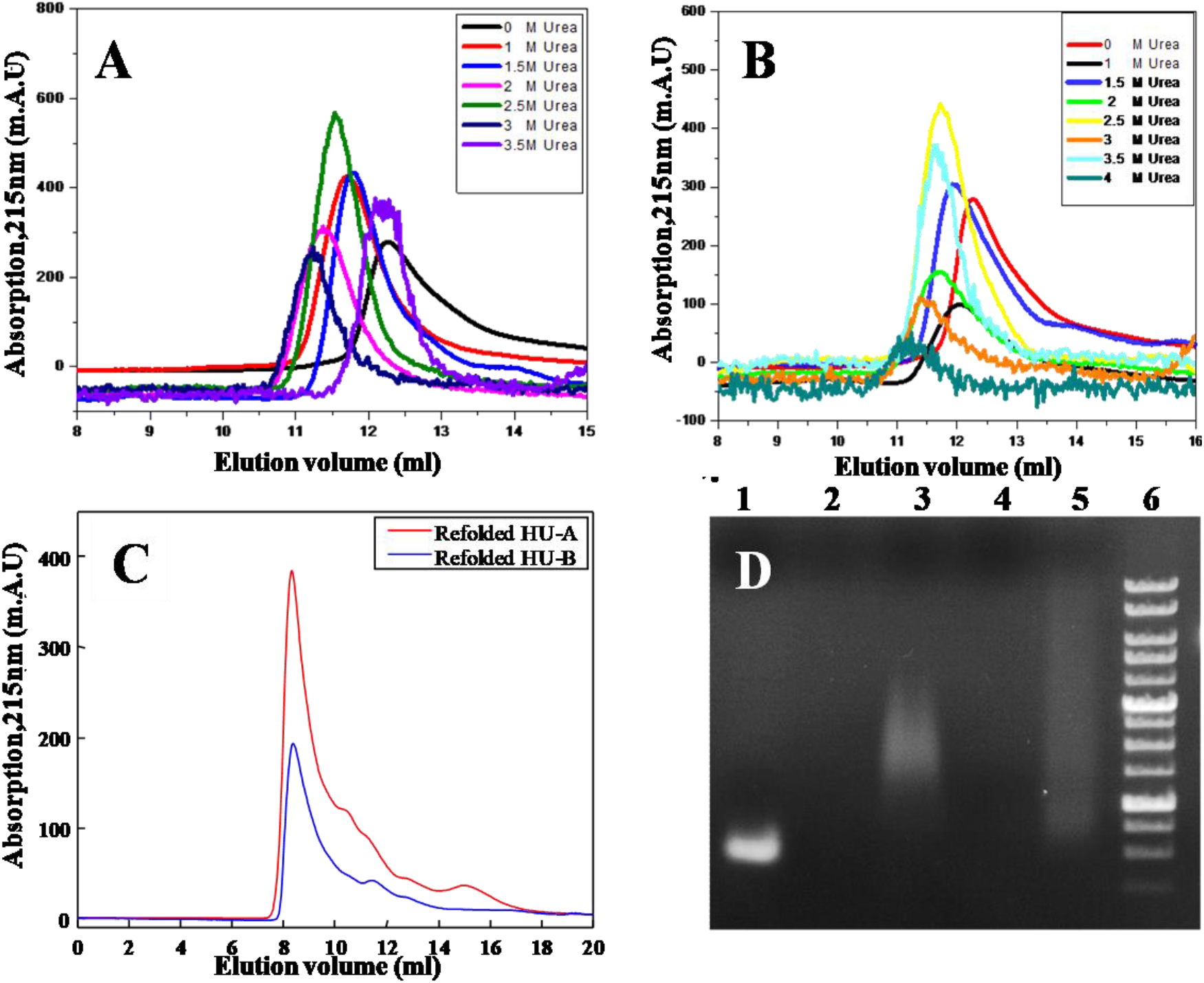
Behavior of chemically unfolded and refolded histidine-tagged HU. *Panel A.* HU-A incubated in urea and subjected to gel filtration chromatography at the same urea concentration used for incubation. *Panel B.* HU-B incubated in urea and subjected to gel filtration chromatography at the same urea concentration used for incubation. *Panel C.* Gel filtration chromatographic behavior of HU-A (in red) and HU-B (in blue) after refolding from 6 M urea. *Panel D.* Electrophoretic mobility shift assays of refolded HU electrophoresed with, or without, DNA, using an agarose (0.5 %) gel: Control 4WJ DNA (lane 1), refolded HU-A (lane 2), refolded HU-A with 4WJ DNA (lane 3), refolded HU-B (lane 4), refolded HU-B with 4WJ DNA (lane 5), 100 BP resolution DNA molecular weight markers ladder (lane 6).

#### Gel-filtration chromatography reveals that refolding from chemically-denatured state gives rise to DNA binding-competent soluble aggregates of HU-A and HU-B

Figure 5C shows the gel filtration behavior of HU-A and HU-B exposed overnight to 6 M urea and dialyzed to allow complete removal of denaturant. The figure shows that both homologs form large soluble aggregates that elute in the void volume of the gel filtration column, suggesting that refolding does not lead to formation of dimers. However, the electrophoretic mobility shift assay data in Figure 5D establishes that these soluble aggregates of HU-A and HU-B are DNA binding-competent, because they slow down the migration of 4WJ DNA.

#### Glutaraldehyde crosslinking reveals that the persistent sub-structure resisting unfolding and dissociation involves the dimeric subunit interface

To address the question of whether there is sufficient survival of some core structure in HU-A and HU-B which is associated with inter-subunit interactions, and the retention of dimeric, or tetrameric, states at high denaturant concentrations, we performed glutaradehyde crosslinking experiments using protein samples that had previously been incubated overnight in the presence of different concentrations of urea. The results are shown in Figures 6A and 6B for HU-B, and 6C for HU-A. From previous examination of Figure 4C, it is evident that the C_m_ value of HU-B with urea is 2.14 M. Figures 6A and 6B show that dimeric species survive in HU-B samples much above the urea concentration of 2.14 M, even at 4 M urea, with even tetrameric species surviving at 2.5 M urea which is above the urea concentration of 2.14 M. It may be noted that 4 M urea is five-sixths of the way to complete unfolding of HU-B. This suggests that HU-B undergoes non-two-state unfolding with subunit dissociation uncoupled from the molecule’s unfolding, since there is clear persistence of sufficient structure in the ‘substantially-unfolded’ state to facilitate retention of dimeric or tetrameric state(s) of HU. An entirely similar result is obtained with HU-A, as seen in Figure 6C, in which dimeric species are observed even at 2.5 M urea, despite HU-A displaying a C_m_ of only 1.77 M urea. Therefore, in both homologs, glutaraldehyde manages to crosslink monomers (suggesting persistence of dimers) at urea concentrations that are significantly in excess of the C_m_ values for urea.

**Figure 6:**
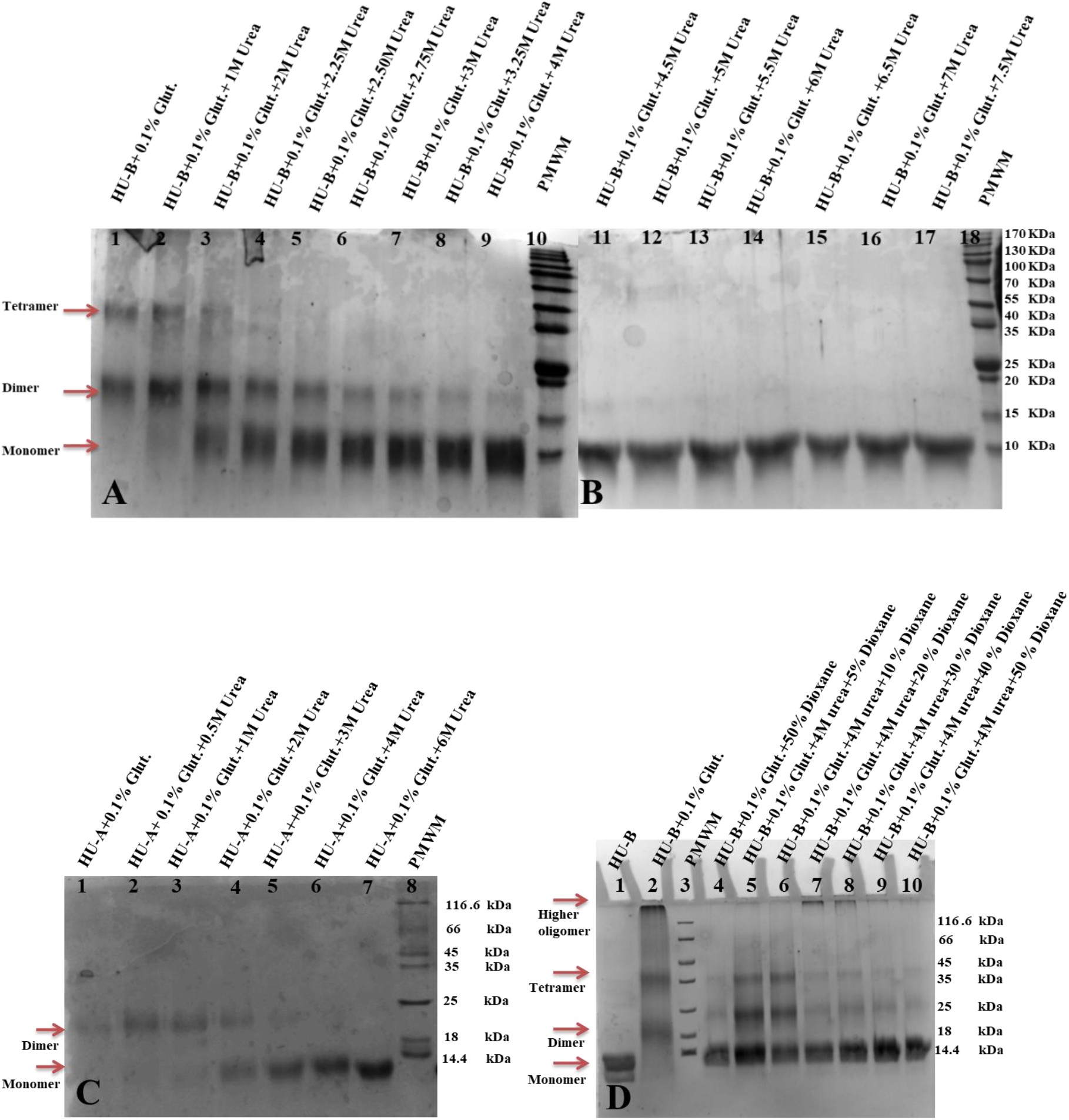
Glutaraldehyde crosslinking studies to estimate the strength of the dimeric interface of HU. *Panel A.* HU-B crosslinking behavior at urea concentrations ≤ 4 M urea. *Panel B.* HU-B crosslinking behavior at urea concentrations ≥ 4 M urea. *Panel C.* HU-A crosslinking behavior at urea concentrations between 0 and 6 M. *Panel D.* HU-B crosslinking behavior in the presence of 1,4-dioxane alone, or in the presence of 1,4-dioxane and urea. Details of glutaraldehyde concentrations used, are mentioned above each lane. PMWM stands for protein molecular weight markers.

#### A hydrophobic solvent (dioxane) disrupts crosslinking, suggesting that hydrophobic interactions at the subunit interface underlie the persistent sub-structure

The bulk of the structures of HU-A, or HU-B, is stabilized by hydrogen bonding. The entire NTD is stabilized by helices that are susceptible to undergoing unfolding through a helix-coil transition upon disruption of hydrogen bonding by urea. On the other hand, significant portions of the CTD are stabilized by a combination of hydrogen-bonding and hydrophobic interactions (notwithstanding the IDR regions which undergo electrostatic interactions with DNA to achieve structure; otherwise remaining unstructured). We decided to examine whether a non-polar, water-miscible, solvent such as 1,4-dioxane could be used to effect complete dissociation of HU-B in the presence of urea. Lanes 5-10 in Figure 6D demonstrate that use of 1,4-dioxane reduces the survival of dimeric and tetrameric species in HU-B in a dose-dependent manner, in the presence of 4 M urea. Further, Lane 4 demonstrates that 50 % dioxane is able to substantially obviate the survival of dimeric and tetrameric species even without the presence of urea, to the same degree that is seen with use of 50 % dioxane in the presence of 4 M urea. This establishes that the dimeric interface holds the key to the structural stability of HU, with destruction of this interface leading to unfolding and dissociation. Further, Supporting Information Figure S2 presents data for HU-B refolded through dialysis of urea and dioxane, or dioxane alone. The figure shows that refolding converts most of the protein into dimeric and tetrameric form (with glutaraldehyde binding reducing staining by Coomassie, as known)^23^ in both situations.

### Stability of HU dimers to thermal unfolding also reveals a persistent hydrophobically-stabilized dimeric interface in histidine-tagged HU that could frustrate subunit exchange

#### Circular dichroism reveals limited (partial) thermal unfolding of HU-A and HU-B, with hints of persistence of sub-structure due to hydrophobic interactions stabilized further by heat

The relative abilities of the three-dimensional structures of HU-A and HU-B to withstand the application of heat was assessed by spectroscopic means. CD spectra were collected at temperatures between 20 °C and 90 °C, during heating of the protein at a controlled rate (2 °C/min) and also during cooling of the protein from 90 °C to 20 °C at the same rate. Figures 7A and 7B, respectively, plot the changes in the mean residue ellipticity (MRE) at 222 nm, for HU-A and HU-B, during heating (red curve) and cooling (blue curve). From the data, it is clear that the magnitude of reduction in helix content with temperature, as well as the temperature eliciting 50 % reduction in CD signal strength at 222 nm (T_m_), happen to be different for HU-A, and HU-B. With HU-A, the T_m_ ~ 40.6 °C, whereas for HU-B, the T_m_ is ~ 51.2 °C, establishing that HU-A is less stable to thermal unfolding than HU-B, exactly as was seen to be the case for unfolding by chemical denaturants, as already noted above. Further, it is evident from the spectra that structural changes in HU-A are initiated upon heating from 20 °C, whereas in HU-B no changes are seen below 30 °C. Figures 7A and 7B also support the observation that HU-B is more structurally stable than HU-A, since it is evident from these figures that, at 90 °C, there is more residual structure in HU-B, than in HU-A. Further, it must be noted that the amount of residual structure persisting after thermal unfolding has been allowed to saturate is much greater than the residual structure seen after chemical unfolding in the presence of 6 M Gdm.HCl, or 8 M urea. This difference supports the likely importance of hydrophobic interactions to the persistence of a stable sub-structure in HU that is resistant to unfolding, since the strength of hydrophobic interactions increases with increasing temperature in the range of 40-140 °C, in aqueous solutions, and this would cause it to interfere to a greater extent with thermal unfolding than chemical unfolding. Indeed, as shown in Supporting Information Figure S3, glutaraldehyde crosslinking performed at 90 °C reveals the persistence of dimeric and tetrameric forms of HU even at this temperature.

**Figure 7:**
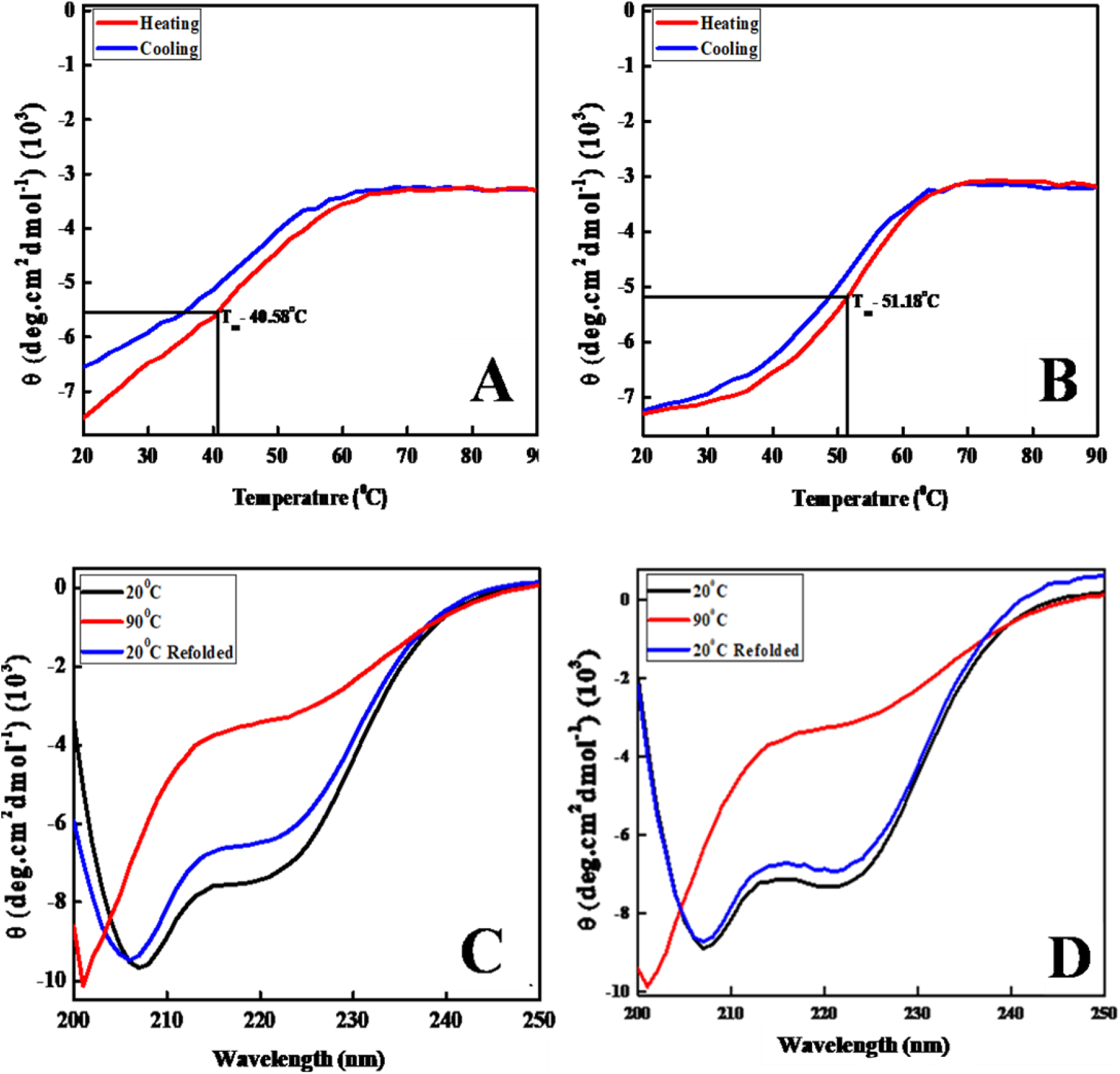
CD spectral mean residue ellipticity (MRE) data monitoring thermal unfolding and refolding of histidine-tagged HU-A and HU-B. *Panel A.* Changes in MRE of HU-A at 222 nm during thermal unfolding (red) and refolding from a thermally-unfolded state (blue). *Panel B.* Changes in MRE of HU-B at 222 nm during thermal unfolding (red) and refolding from a thermally-unfolded state (blue). *Panel C.* Far-UV CD spectra of HU-A, before heating (black), in the heated state (red), and after cooling (blue). *Panel D.* Far-UV CD spectra of HU-B, before heating (black), in the heated state (red), and after cooling (blue).

#### Refolding of HU-A and HU-B occurs from partially thermally-unfolded state(s), possibly owing to retention of a persistent sub-structure

Figures 7C and 7D show the CD spectra of HU-A, and HU-B, respectively, collected at 20 °C, i.e., before heating (black curve), then at 90 °C in the heated state (red curve), and once again at 20 °C after cooling (blue curve). The data establishes that there is some hysteresis evident between the heating (unfolding) and cooling (refolding) curves, with HU-B returning to nearly the same secondary structural content after cooling, but with HU-A displaying significantly less negative MRE at 222 nm after cooling, suggesting a somewhat poorer refolding efficiency than HU-B. The entire spectra collected before heating, and after cooling, which are shown in Figures 7C and 7D also support the occurrence of this hysteresis, in terms of there being a larger difference between initial and final states in HU-A than in HU-B.

#### Reassociation of refolded HU-A and HU-B into DNA binding-competent dimers from a partially thermally-unfolded state, possibly owing to retention of the persistent sub-structure

We then examined the protein refolded from a thermally-unfolded state to verify whether it had formed predominantly dimeric HU. Figures 8A and 8B establish that the gel filtration chromatographic elution profiles of unheated HU-A and HU-B, and their heated-and-cooled forms, overlap sufficiently for one to conclude that both homologs are capable of refolding to dimeric state. Figure 8C shows that these dimers obtained through refolding of partially thermally-unfolded HU (from 90 °C) are capable of binding to 4WJ DNA and displaying electrophoretic mobility shift.

**Figure 8:**
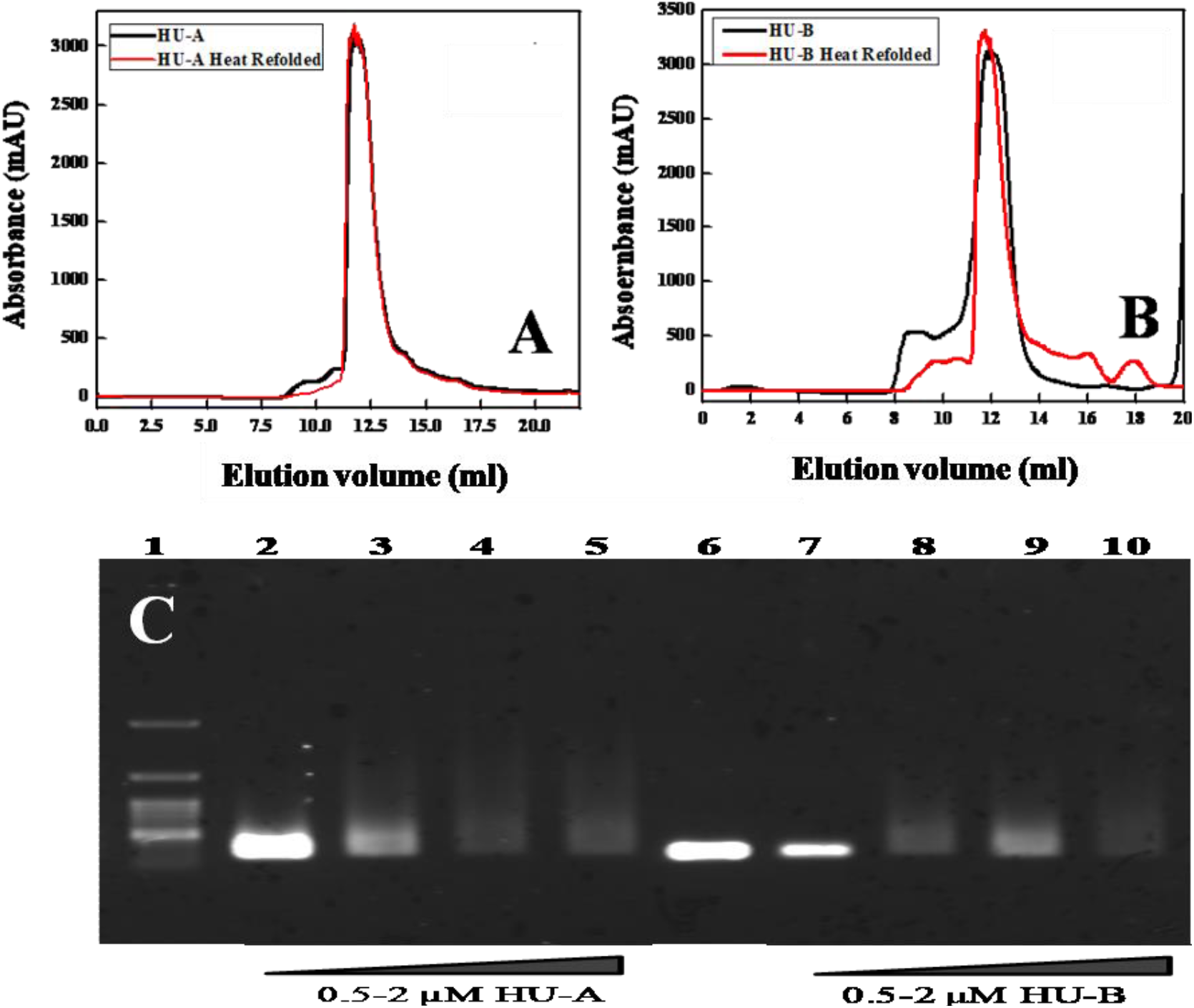
Behavior of histidine-tagged HU before and after thermal unfolding. *Panel A.* Gel filtration elution profile of unheated HU-A (in black) and HU-A refolded after exposure to 90 °C (in red). *Panel B.* Gel filtration elution profile of unheated HU-B (in black) and HU-B refolded after exposure to 90 °C (in red). *Panel C.* Electrophoretic mobility shift assays of HU-A and HU-B following refolding after thermal unfolding, through electrophoresis with DNA, using an agarose (0.5 %) gel: 100 BP resolution DNA molecular weigh markers ladder (lane 1); control 2 μM 4WJ DNA alone (lane 6); 2 μM 4WJ DNA with refolded HU-A [0.5 μM (lane 2); 1.0 μM (lane 3); 1.5 μM (lane 4); 2.0 μM (lane 5)]; and 2 μM 4WJ DNA with refolded HU-B [0.5 μM (lane 7); 1.0 μM (lane 8); 1.5 μM (lane 9); 2.0 μM (lane 10)].

## DISCUSSION

In *E. coli* cells, HU exists as two homologous isoforms, HU-A, and HU-B. These proteins, which are produced from the genes, *hupA* and *hupB*, are reported to vary in amount and relative abundance with the stage of growth of a population. As already mentioned in the introductory section, HU-AA homo-dimers are thought to exist mainly during the early log phase, while HU-BB homodimers are thought to exist mainly in the early and mid-exponential phase, and HU-AB heterodimers are thought to exist mainly during the late exponential phase and stationary phase.^6^ In the stationary phase, heterodimers are thought to be the predominant population.^5^ The formation of the heterodimer has been shown to alter the extent of HU-mediated compaction of DNA, by changing HU oligomerization status.^13^ The oligomeric status of the HU-AB hetero-dimer, in terms of the fraction of any HU population that forms higher-order associations over and above the dimeric state, has been reported to be intermediate to that of HU-AA (which is predominantly dimeric) and HU-BB (which exists as dimers, tetramers, hexamers and octamers). It may be conjectured that heterodimers are preferentially-formed to facilitate the higher compaction of bacterial nucleoids during the stationary phase.^24^ In the existing literature, the mechanism discussed for formation of HU-AB hetero-dimers involves exchange of subunits between HU-AA homo-dimers and HU-BB homo-dimers.^5,6,20^ In principle, of course, HU hetero-dimers could only conceivably form, through the exchange of subunits between HU homo-dimers, if facile dissociation of HU monomer-monomer interfaces can be conceived to occur feasibly within HU dimers, during their dissociation (potentially associated with a certain amount of chain reorganization, due to the nature of the dimeric interface, which involves intimate NTD-NTD and CTD-CTD inter-subunit interactions).

Analyses of the structure of the HU dimer, and inter-subunit chain-chain interactions indicates that any problem in the dissociation of subunits (whether in HU-AA, HU-BB, or HU-AB) would be more likely to be caused by the C-terminal half of each chain, i.e., by the CTD, than by the NTD. The NTDs of both chains are constituted entirely of alpha helices anticipated to unfold and dissociate readily through facile helix-coil transitions; however, the CTDs engage in strong hydrophobic interactions between equivalent anti-parallel beta strands contributed by both chains that could potentially survive destabilization of other hydrogen bonding and electrostatic interactions. Wild-type HU is thought to exist predominantly as a heterodimer (to the extent of ~ 90 % of the population) when there are both HU-A and HU-B chains present, suggesting that despite the strong CTD-CTD contacts, subunit exchange occurs in a facile manner. However, earlier reports suggest that N-terminally histidine-tagged HU is purified in the form of homodimers. In the introductory section, we argued that the purification of homodimers of histidine-tagged HU suggests that there could be a subtle phenotype of the protein resulting from its N-terminal tagging; a phenotype manifesting as a problem in dissociation and subunit exchange to generate heterodimers, due to the N-terminus being modified through its extension, and displacement, in the form of the placement of a histidine tag. In HU chains, the third amino acid from the N-terminus is a lysine, and it is conceivable that the charge-charge repulsions between the N-terminal alpha amino group and the epsilon amino group of residue K3 destabilizes the CTD-CTD contact and promotes facile subunit exchange, whereas removal of the N-terminal charge and its placement several amino acids away reduces this destabilizing influence, and allows the CTD-CTD interactions to become stronger, and manifest as a reluctance to dissociate and allow subunit exchange.

Thus, we resolved to first confirm the observation for ourselves that histidine-tagged HU is indeed homodimeric. For this, we used mass spectrometry. We confirmed that it is indeed homodimeric, with no trace of the chain of the other identity. Then we examined whether histidine-tagged HU has a lower propensity to heterodimerize, through the swapping of subunits upon mixing, or through *de novo* assembly during coexpression. We confirmed that less than 5 % heterodimers are formed, through FRET and *in vivo* coexpression of HU-A and HU-B chains. We then resolved to examine the unfolding and dissociation of HU-A and HU-B by a combination of spectroscopic, chromatographic and electrophoretic methods, to explore whether the subunits do indeed fail to dissociate in a facile manner, to shed light on the low extent of heterodimer formation in comparison with non-histidine-tagged forms of HU which heterodimerize to the extent of ~ 90 %. We confirmed that histidine-tagged HU indeed cannot heterodimerize in a facile manner. During chemical denaturation, the HU dimer initially unfolds into a substantially-unfolded but nominally-dimeric state with a greater hydrodynamic volume, residual secondary structure, and potential for inter-chain crosslinking by glutaraldehyde. We found that HU-B exhibits these tendencies to a greater degree than HU-A. Urea unfolding studies elicited these differences to a greater degree than Gdm.HCl. After first unfolding into a partially-unfolded dimer, HU dissociates into unfolded monomers, with such dissociation facilitated by the non-polar solvent dioxane, supporting the probability that the dimeric interface that displays a tendency to persist/survive involves hydrophobic interactions between the two CTDs of the two chains; a possibility that is also supported by the fact that the helical signal (from the NTDs) is almost completely destroyed in the CD spectra, while residual signal attributable to beta sheets remains, even in a state that is susceptible to crosslinking, and which shows progressively larger hydrodynamic volume than a folded monomer during exposure to higher concentrations of urea. Figure 9 presents a summary view of our conclusions regarding the unfolding and dissociation of N-terminally histidine-tagged HU-AA and HU-BB. In the figure, the original N-terminus is shown (without a schematic of the histidine tag) to emphasize that the N-terminus is ordinarily snugly packed against a region of the protein not distant from the CTD-CTD interaction characterized by hydrophobic interactions between antiparallel beta sheets.

**Figure 9:**
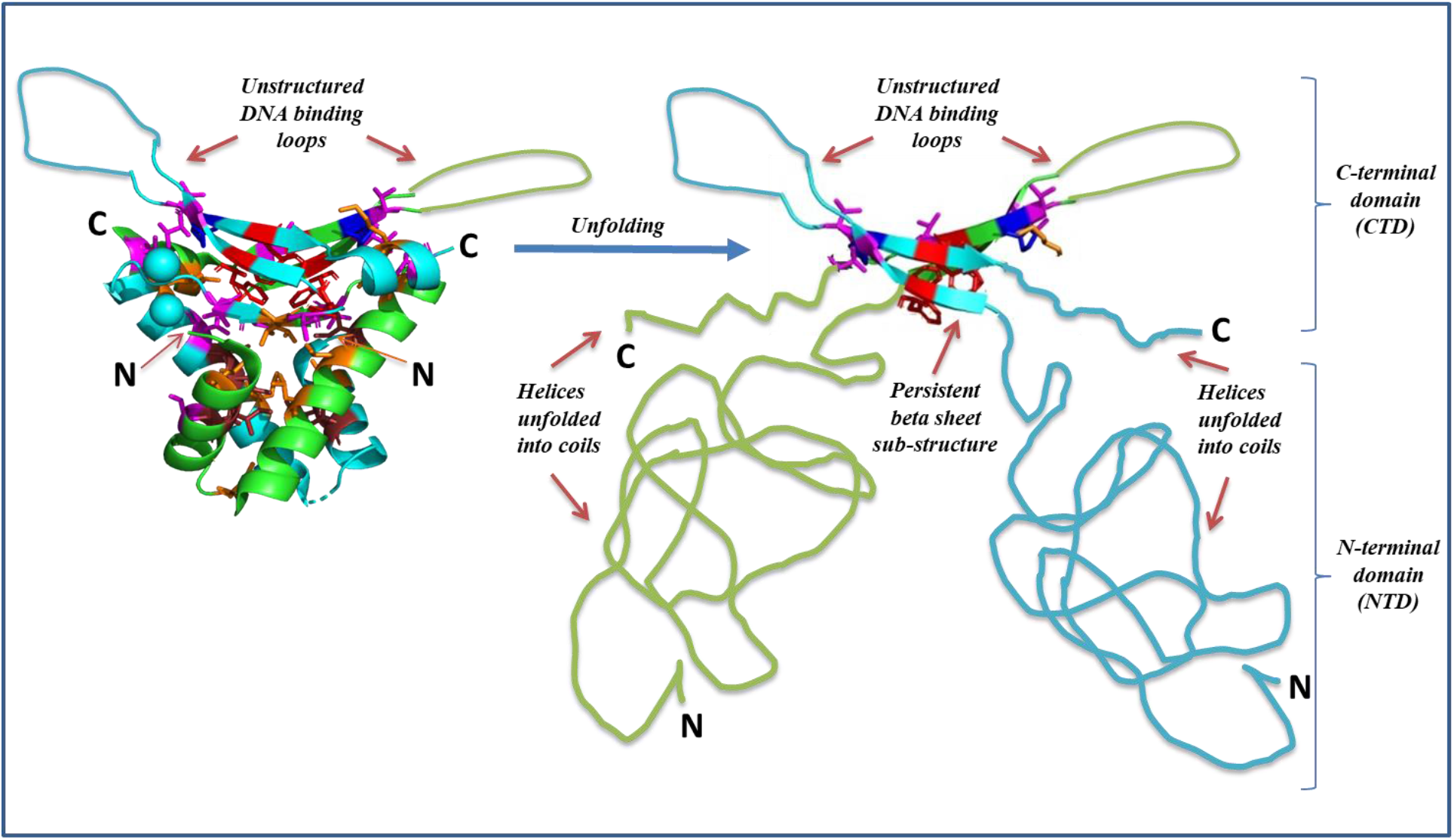
Proposed mechanism of HU unfolding and dissociation. *Panel A.* Ribbon diagram of the HU-AB heterodimer (PDB ID:2O97), shown here as a representative of the generic HU homodimer, with positions of the N- and C-termini marked, and showing all inter-subunit NTD-NTD and CTD-CTD interactions, as well as hydrophobic residue side chains (shown using a stick model). The region marked IDR represents a schematic of the unstructured DNA binding loop that is flexible in the absence of DNA. *Panel B.* A schematic of the proposed substantially-unfolded form of HU that, however, remains nominally dimeric due to persistence of inter-subunit CTD-CTD interactions between antiparallel beta sheets engaged in hydrophobic interactions, prior to any complete unfolding and dissociation.

In regard of each of these behaviors, HU-B could be shown to display them to a much greater extent that HU-A. However, we are not certain of the reason for this difference, which could lie in the sequence, or the structure. The sequences of the HU-A and HU-B polypeptide chains are shown in Supporting Information Figure S4, which shows both an alignment of the two sequences and also the secondary structures adopted by different stretches of the sequences, in differently colored fonts. The thermal denaturation data is in line with the chemical denaturation data with some differences, including the greater retention of residual structure. Such greater retention could be explained by the fact that hydrophobic interactions tend to be strengthened with increasing temperature, such that if hydrophobic interactions were predominantly involved in the stabilization of a beta sheet-based structure, this could explain the survival of beta sheets in the CTDs of HU with heating.

The likelihood of homo-dimers or hetero-dimers forming on the basis of encounters between chains in a nascent stage of folding, rather than exchange of subunits between fully-folded and associated chains, is clearly higher for histidine-tagged HU than for non-histidine-tagged HU. Thus, the homo-dimers or hetero-dimers that are once formed have long lifetimes, due to inter-subunit hydrophobic contacts. Any hetero-dimers formed by histidine-tagged HU must form more through *de novo* folding and association of chains, than through exchange of subunits. The observations further confound our understanding of HU, by demonstrating that the lack of heterodimer formation is not essential for *E. coli* growth. Why then does *E. coli* have two homologs with such an interestingly coordinated expression of homologs? The answer probably lies in the behavior of *E. coli* in the colon, rather than in the laboratory.

## Protein Accession Numbers

The UNIPROT Accession Number for the protein referred to as [HU-A](https://www.uniprot.org/) is P0ACF0, and for the protein referred to as [HU-B](https://www.uniprot.org/) is P0ACF4.

## Acknowledgements

KA thanks the University grant commission, Government of India, for a doctoral fellowship. BT thanks the Council of Scientific and Industrial Research (CSIR), New Delhi, for a doctoral fellowship. AM thanks the Department of biotechnology, Government of India, for a doctoral fellowship. PG thanks the Ministry of Human Resource Development (MHRD), Government of India, for a Centre of Excellence Grant (MHRD-14-0064) in Protein Science, Design and Engineering.

## SUPPORTING INFORMATION FOR MANUSCRIPT

**Supporting Information Figure S1:**
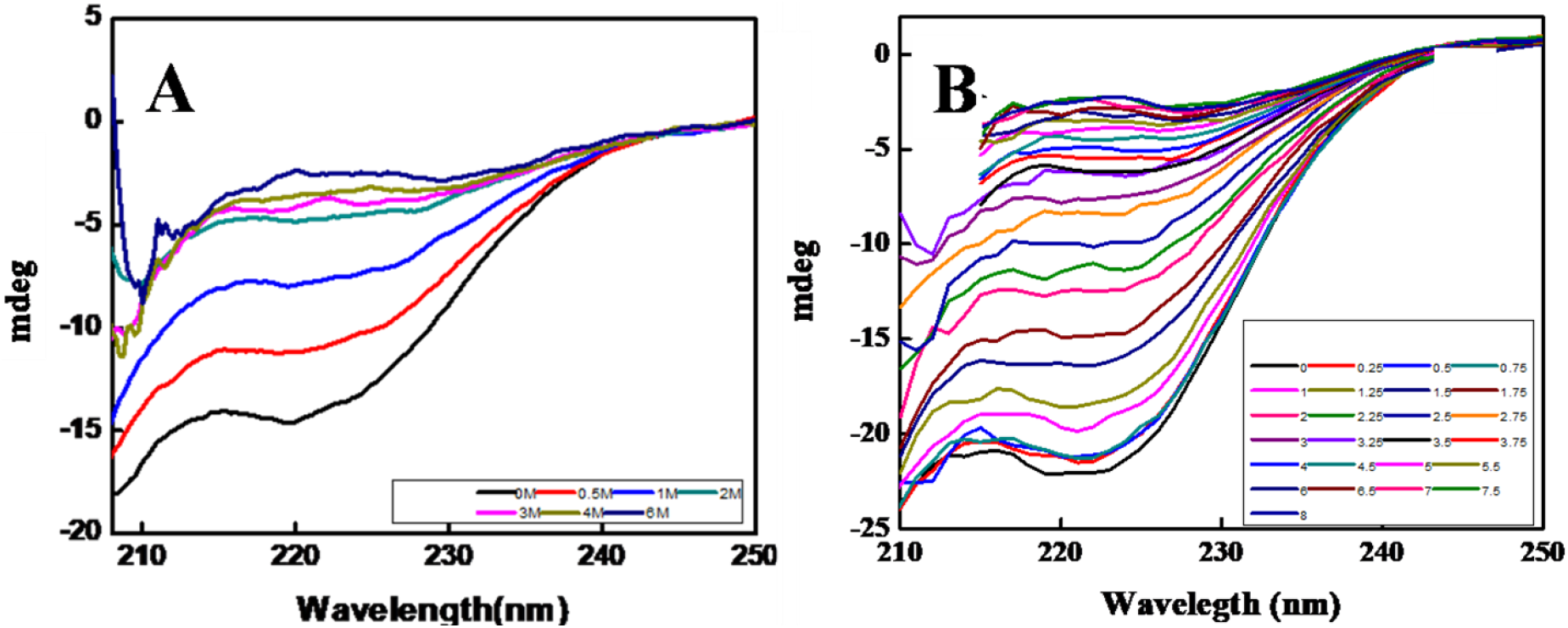
Secondary structure changes in HU-A and HU-B upon incubation in different molar concentration of urea monitored using circular dichorism spectroscopy. *Panel A.* Secondary structure changes for HU-A. *Panel B.* Secondary structure changes for HU-B.

**Supporting Information Figure S2:**
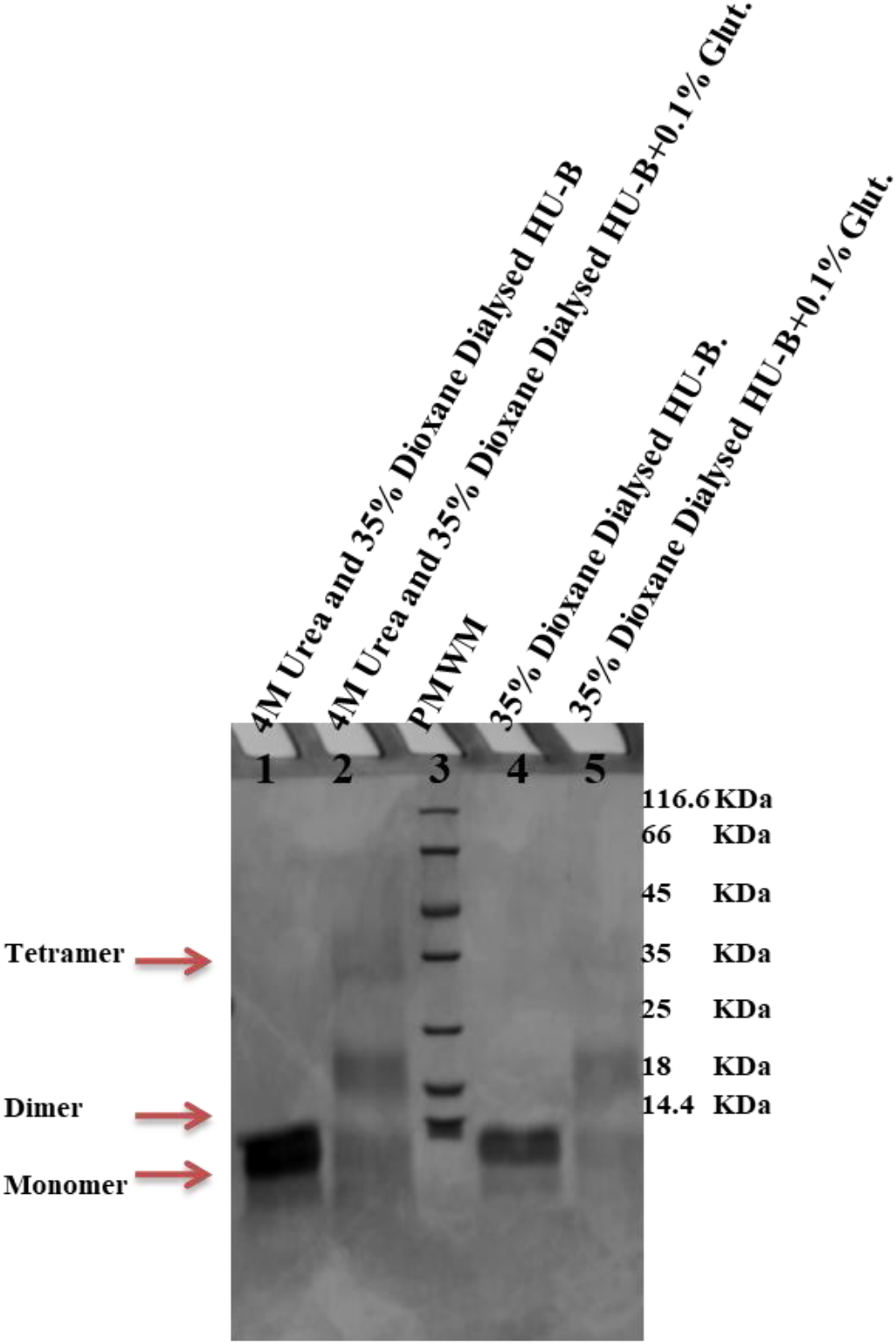
Glutaraldehyde crosslinking to assess refolding of HU into dimer from urea and/or 1,4-dioxane. HU-B crosslinked after removal of 1,4-dioxane alone, or both 1,4-dioxane and urea. All details, including urea and glutaraldehyde concentrations used, are mentioned above each lane. PMWM stands for protein molecular weight markers.

**Supporting Information Figure S3:**
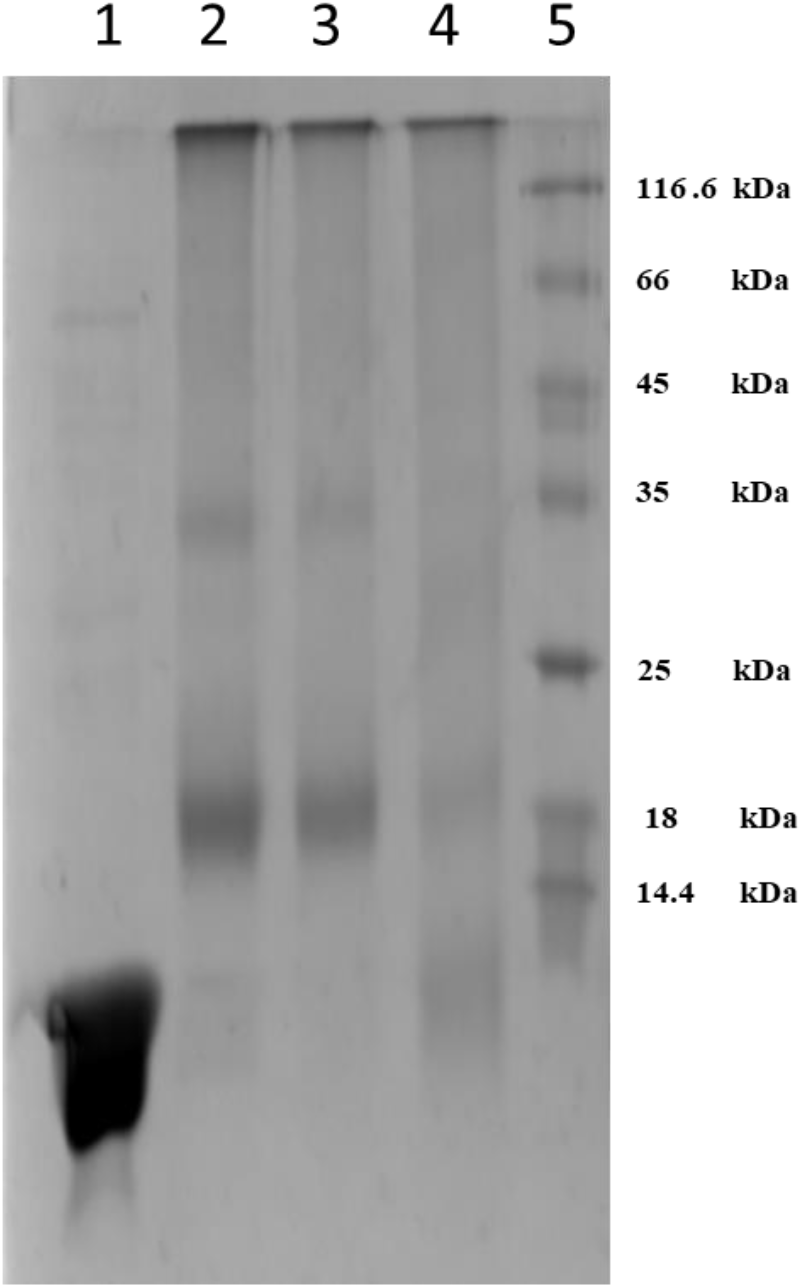
Glutaraldehyde crosslinking to assess persistence of dimeric structure in histidine-tagged HU-B before and after thermal unfolding (at 90 deg C). Lane 1 shows histidine-tagged HU-B. Lane 2 shows HU-B crosslinked with 0.1 % glutaraldehyde before heating, at room temperature. Lane 3 shows HU-B crosslinked with 0.2 % glutaraldehyde, before heating, at room temperature. Lane 4 shows HU-B heated at 90 deg C, and crosslinked with 0.1 % glutaraldehyde at 90 deg C. Lane 5 shows protein molecular weight markers.

**Supporting Information Figure S4:**
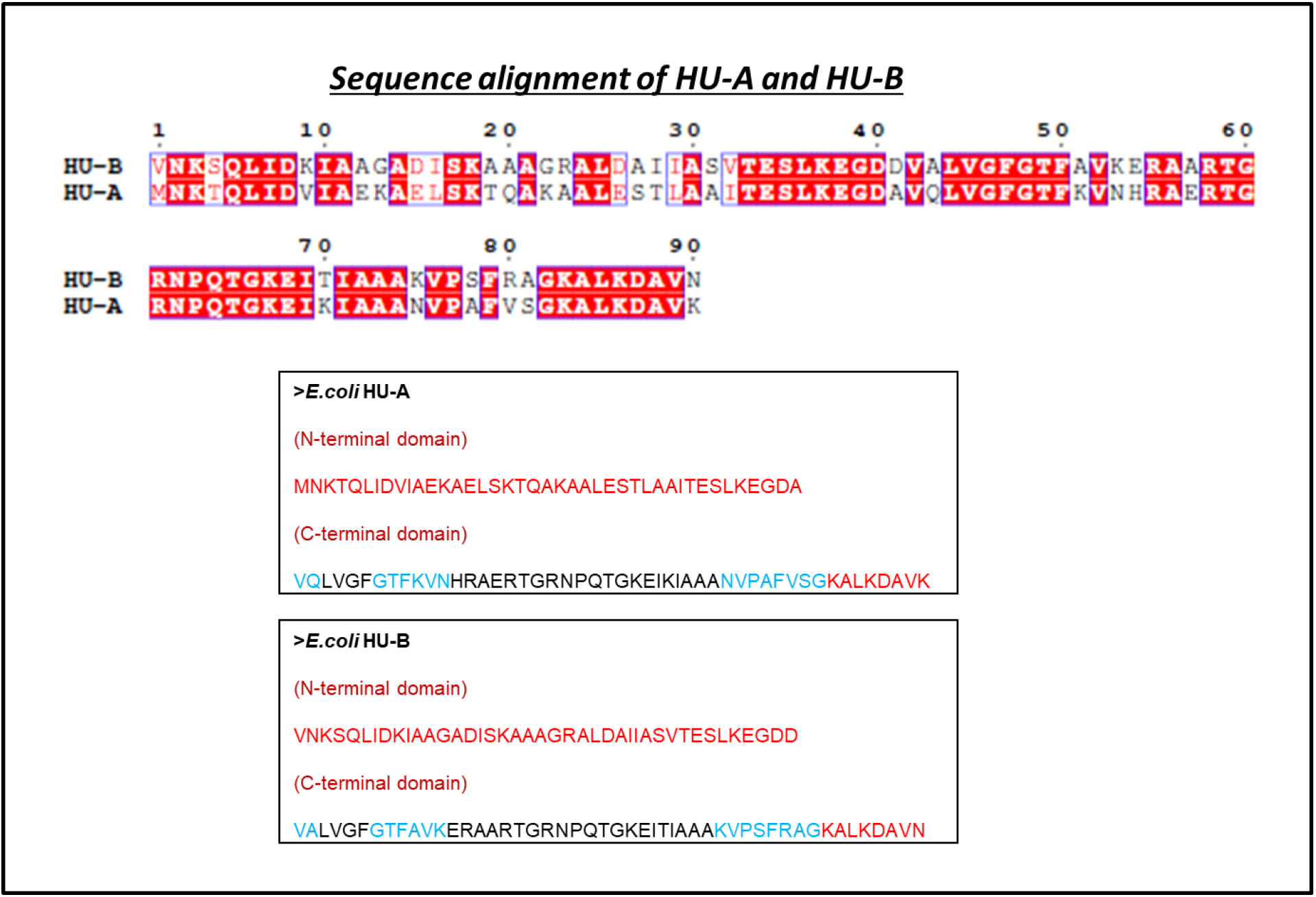
Sequences of wild-type HU-A and HU-B lacking histidine tags. *Top panel.* Sequence alignment of HU-A and HU-B. *Middle panel.* Sequence of HU-A showing the N-terminal and C-terminal domains separately, with residues in helical structures shown in red, residues in sheet structures shown in blue, and residues in intrinsically disordered regions (IDRs) including the main DNA-binding loops, shown in black. *Bottom panel.* Sequence of HU-B showing the N-terminal and C-terminal domains separately, with residues in helical structures shown in red, residues in sheet structures shown in blue, and residues in intrinsically disordered regions (IDRs) including the main DNA-binding loops, shown in black.

